# Quantitative agent-based modeling reveals mechanical stress response of growing tumor spheroids is predictable over various growth conditions and cell lines

**DOI:** 10.1101/122614

**Authors:** Paul Van Liedekerke, Johannes Neitsch, Tim Johann, Kevin Alessandri, Pierre Nassoy, Dirk Drasdo

**Affiliations:** INRIA de Paris and Sorbonne Universités UPMC Univ paris 6, LJLL, France; IZBI, University of Leipzig, Hartelstrasse 16-18, Leipzig, Germany; Institut d’Optique Graduate School, Talence, France; IfADo - Leibniz Research Centre for Working Environment and Human Factors, Ardeystrasse 67, Dortmund, Germany

## Abstract

Model simulations indicate that the response of growing cell populations on mechanical stress follows the same functional relationship and is predictable over different cell lines and growth conditions despite the response curves look largely different. We develop a hybrid model strategy in which cells are represented by coarse-grained individual units calibrated with a high resolution cell model and parameterized measurable biophysical and cell-biological parameters. Cell cycle progression in our model is controlled by volumetric strain, the latter being derived from a bio-mechanical relation between applied pressure and cell compressibility. After parameter calibration from experiments with mouse colon carcinoma cells growing against the resistance of an elastic alginate capsule, the model adequately predicts the growth curve in i) soft and rigid capsules, ii) in different experimental conditions where the mechanical stress is generated by osmosis via a high molecular weight dextran solution, and iii) for other cell types with varying doubling times. Our model simulation results suggest that the growth response of cell population upon externally applied mechanical stress is the same, as it can be quantitatively predicted using the same growth progression function.

**Author summary:** The effect of mechanical resistance on the growth of tumor cells remains today largely unquantified. We studied data from two different experimental setups that monitor the growth of tumor cells under mechanical compression. The existing data in the first experiment examined growing CT26 cells in an elastic permeable capsule. In the second experiment, growth of tumor cells under osmotic stress of the same cell line as well as other cell lines were studied. We have developed and agent-based model with measurable biophysical and cell-biological parameters that can simulate both experiments. Cell cycle progression in our model is a Hill function of cell volumetric strain, derived from a bio-mechanical relation between applied pressure and cell compressibility. After calibration of the model parameters within the data of the first experiment, we are able predict the growth rates in the second experiment. We show that that the growth response of cell populations upon externally applied mechanical stress in the two different experiments and over different cell lines can be predicted using the same growth progression function once the growth kinetics of the cell lines in abscence of mechanical stress is known.

## 1 Introduction

Mechanotransduction is the mechanism by which cells transform an external mechanical stimulus into internal signals. It emerges in many cellular processes, such as embryonic development and tumor growth [1]. Cell growth in a confined environment such as provided by the stroma and surrounding tissues increases cell density and affects the balance between cell proliferation and death in tissue homeostasis [2,3]. Tumor spheroids have long been considered as appropriate in vitro models for tumors [4]. While the dynamics of freely growing spheroids has been extensively studied both experimentally [5] and numerically (e.g. [6,7]), more recent experiments have also addressed the growth of spheroids under mechanical stress.

Helmlinger et al. (1997) and later Cheng et al. (2009) and Mills et al. (2014) [8–10] experimentally investigated the growth of spheroids embedded in agarose gel pads at varying agarose concentration as a tunable parameter for the stiffness of the surrounding medium. Other approaches such as the application of an osmotic pressure determined by a dextran polymer solution have also been developed to investigate the impact of external pressure on spheroid growth [11]. In all cases mechanical stress was reported to slow down or inhibit spheroid growth. Delarue et al. [12] suggested that growth stagnation is related to a volume decrease of the cells. However, a quantitative relation between pressure and cell fate is not reached yet. The works of Helmlinger et al. [8] and their follow-ups have inspired a number of theoretical papers aiming at explaining the observations, either based on continuum approaches considering locally averaged variables (e.g. for density and momentum, for overview see [13]) [3,14–17], or by agent-based models (ABMs) representing each individual cell [18,19] belonging to the class of models, which are extended and refined in the presented work. For example, the growth kinetics of multicellular spheroids (MCS) embedded in agarose gel as observed by Helmlinger et al. [8] could be largely reproduced, if cell cycle progression was assumed to be inhibited either above a certain threshold pressure or below a certain threshold distance between the cell centers, whereby growth inhibition occurred at different spheroid sizes for different densities of extracellular material [18]. However, the model developed in that reference has no notion of cell shape, hence does not permit definition of cell volume, thus pressure and compression cannot be physically correctly related [20].

Here, we first establish a computational model to quantitatively explain the growth kinetics and patterns found for CT26 (mouse colon carcinoma cell line) multi-cellular spheroids constrained by a spherical elastic capsule, partially based on data previously published [21] and partially based on new data introduced below. This novel experimental technique, called the “cellular capsule technology” [21] allows to measure the average pressure exerted by the cell aggregate onto the calibrated capsule by monitoring the radial expansion of the shell once confluence is reached. Pressure can be recorded over periods as long as a week and the histological data collected and analyzed on fixed and sliced spheroids can provide snapshots of the spatial multicellular pattern.

We refer to this experimental technique as “Experiment I”. The thickness, and thus the stiffness of the capsule, was varied to mimic different mechanical resistance conditions.

Delarue et al. (2014) [12] investigated the effect of mechanical stress on MCS growth using the same cell line in a different experimental setting. We exploit these results to challenge our model and determine whether the same computational model designed to match experiment I is capable to quantitatively explain also this experiment (referred to as “experiment II”). In experiment II, mechanical compression was imposed using the osmotic effects induced by a dextran solution. The main difference between those two experiments is that whereas the pressure gradually increases with increasing deformation of the elastic capsule in experiment I, in experiment II a constant stress is applied due to osmotic forces in the absence of any obstructing tissue (see Figure 1A).

**Fig 1.**
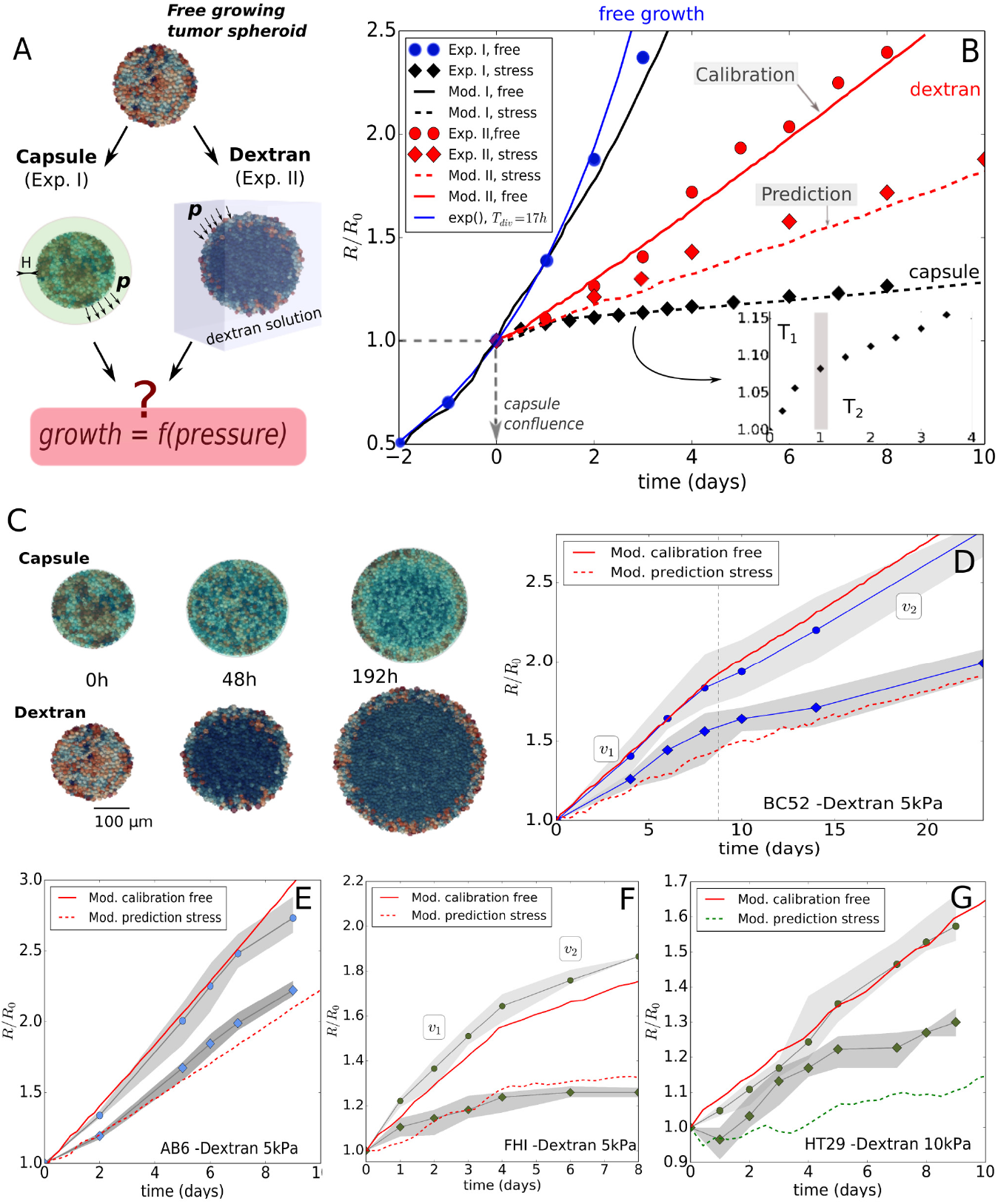
Summary of key experimental and simulation results. (A)Two experiments setups for growing spheroids considered in this study. In experiment I, the spheroid is in mechanical contact with a capsule, and the mechanical resistance is determined by the wall thickness H. In experiment II, the spheroid is immersed in a dextran polymer solution, and the mechanical resistance originates from the osmotic pressure related to the dextran concentration. (B) Radial growth curves data of the spheroids in units of *R*_0_ (= 100 *μ*m), for experiment I and II and respective model runs. The blue full circles are the free growth data for CT26, from [21]. The thin blue line indicates theoretical pure exponential growth with doubling time of 17*h*. The data starts deviating from an exponential after 2 days. The other lines are simulation results. The black dashed line indicates the optimal parameter set for the stress response in experiment I, performed with final model I. The full black line indicates the same model run for free growth in Exp.I. After re-calibration of one model parameter in model I for the Exp.II conditions in absence of dextran (full red line), the model (referred to as model II to stress the change of the parameter) predicts the stress response in experiment II (red dashed line). (C) Simulation snapshots of both experiments. The cells are colored according to their volume (cells at the border are larger than in the interior). (D-G) Model simulations for Exp.II for the cell lines BC52, AB6, FHI and HT29, respectively. Full red lines represent the same initial calibration procedure, while red dashed lines represent the predicted stress conditions. The stress conditions are p = 5 kPa for AB6, FHI and BC52, and *p* = 10kPa for HT29 (see section Validation of model for experiment II: other cell lines).

In this paper, we aim to decipher and quantify certain mechanisms of spheroid growth altered by mechanical stress. At this stage, we establish a robust computational approach that can be applied to various systems (cell lines and experimental procedures) and that allows to recapitulate the growth dynamics and the observed cellular patterns. We will show that this can be reached with a minimal number of hypotheses without having to explicitly integrate specific molecular pathways. Gaining insight in the molecular mechanisms would require additional challenging experiments in which the pathways are selectively inhibited or enhanced in a three-dimensional environment, and would add further parameters to the model. To the best of our knowledge, a specific mechanotransduction molecular pathway has been highlighted once, demonstrating the impact of cell volume change on the expression of the proliferation inhibitor *p*27^*Kip*1^ [12].

As modeling technique we here developed an agent-based model. Simulations with ABMs provide a computer experiment representing an idealized version of the true wet-lab experiment [74]. ABMs naturally permit accounting for cell to cell variability and inhomogeneities on small spatial scales as they represent each cell individually. Center-Based Models (CBM) are a prominent representative in the class of ABMs in which forces between cells are calculated as forces between their centers. Center-based models for multicellular systems were derived from conceptual anologies to collodial particle dynamics by re-interpretation of parameters and addition of growth and division processes [48,72]. The model developed here is fully parameterized in terms of physical parameters, which makes each component possible to validate. However, it circumvents difficulties that standard center-based models have at large compression (see [20]) establishing a hybrid modeling strategy to compute the mechanical interaction forces by so-called 3D Deformable Cell Models (DCMs) [67,75]. A DCM displays cell shape explicitly at the expense of high computational cost (see Figure 3). In our hybrid strategy the parameters of the CBM that considers the cell shape only in a statistical, “coarse grained” sense thereby permitting simulations of large cell population sizes, are pre-calibrated from a finer scale DCM. This strategy permits to combine the advantages of the DCM with the short simulation time of the CBM. Both CBM and DCM are parameterized by measurable quantities to identify the possible parameter range of each model parameter and avoid non-physiological parameter choices.

We studied the series of experimental settings in the works [21] and [12] as both utilize a common cell line, and exert stress on growing MCS of that cell line in different experimental settings. The model is then further tested with experiments on other cell lines as provided in the second work.

To unravel the dynamics of MCS subject to external mechanical stress, our modeling strategy is to postulate and implement hypotheses on cell growth, quiescence and death, and iteratively adapt or extend them in case the model simulations are falsified by comparison with the experimental data. Pursuing a similar strategy enabled us to obtain predictions of subsequently validated mechanisms in liver regeneration [22,23]. Based upon analysis of the relation between pressure, cell density and cell compressibility in the two different experiments, our findings suggest that contact inhibition can be regarded as a robust continuous process imposed by a reduction of cell volume as a consequence of increasing pressure and individual cell compressibility. In addition, the high-resolution model shows that potential effects of micro-mechanics at the interface with the capsule may depend on the mechanical properties of the cells.

For the sake of clarity, we below start to first present the minimal model that was able to explain the data, before discussing in which ways simpler models with other hypotheses failed.

## Results

### Experimental observations

*Experiment I:* Following microfluidics-assisted encapsulation of CT26 cells into alginate hollow capsules, the growing aggregates of cells were monitored by phase contrast microscopy (see [21] for details). After the tumor cells reached the inner border of the elastic alginate capsule corresponding to a radius of about 100 *μm* (*t* = 0*d* in Figure 1B), they were observed to further induce a dilatation of the capsule, which is an indicator of the exerted pressure. The capsule expansion was measured from the point of confluence over several days, while histological data of the spheroids were collected at the stage of confluence and at 48h past confluence. Capsules have been designed to generate shells with two different thicknesses. The thin ones (*H*/*R*_0_ ≈ 0.08; *H* = 8*μm*) are the softer while the thick ones (*H*/*R*_0_ ≈ 0.25; *H* = 30*μm*) will mimic a larger mechanical resistance against growth. Besides the data extracted from [21], we have also exploited and analyzed unpublished data corresponding to new sets of experiments in order to critically test the reliability of the method (see Figure 4A). We extract four main observations from these experiments:

(**EI.OI**) In the absence of a capsule, an initial exponential growth stage was observed with doubling time *T_cyc_* = 17*h* [21]. The growth kinetics however starts to deviate from exponential growth for spheroid size (*R* ≈ 175 *μ*m, see Figure 1B).

(**EI.OII**) In the presence of a capsule, the exponential growth is maintained until confluence, i.e. (*R* = *R*_0_ ≈ 100*μ*m), which shows that the capsule is permeable to nutrients and allows normal growth. Once confluence is passed, the time evolution of the capsule radius exhibits two regimes: i) an initial “fast” growth stage *T*_1_(*t* < 1day), crossing over to ii) a “slow” quasi-linear residual growth stage *T*_2_(*t* > 1 day) that at least persists as long as the capsules are monitored, i.e. up to one week. The transition happens roughly at a pressure of ∼ 1.5 kPa, see Figure 4C. The observed long-time growth velocities were ∼ 2 μm/day for the thin capsules (Figure 4A) and 0.7 *μ*m/d for the thick capsules (Figure 5).

(**EI.OIII**) The nuclei density, obtained from cryosections, increases from ∼ 1 nucleus / 100 *μ*m^2^ before confinement, to roughly 2 nuclei / 100 *μ*m^2^ after confluence, with a relatively higher number near the center of the spheroid (1.2 times more compared to the outer regions), and a local increase at the border of the capsule. The distribution and shape of cell nuclei reported in [21] suggests that cells near the capsule border are deformed thus deviating from a spherical shape cells adopt in isolation, while those in the interior look spherically shaped.

(**EI.OIV**) Most of the cells in the core of the spheroid are necrotic after 48*h* of confinement, while the cells located in a peripheral viable rim of roughly two cell layers thickness (λ_*I*_ ≈ 20*μ*m), show viability and proliferative activity during the whole time course of the experiment, including period *T*_2_.

*Experiment II:* in the work of Delarue et al. (2014) [12], CT26 spheroids (initial radius ∼ 100 *μ*m) were grown in a dextran polymer solution. To recover osmotic balance, water expulsion out of the spheroid generates osmotic forces exerted to the outer cells that are transferred as compressive stresses to the interior (bulk) cells. The concentration of dextran regulates the applied pressure.

(**EII.OI**) The growth rate at *p* = 5 kPa is significantly lower than in control spheroids where no pressure is exerted.

(**EII.OII**) The spheroid free growth data does not show an initial exponential phase found in (EI.OI) (Figure 1B). This surprising discrepancy might result from the different culture conditions between both experiments. In experiment I, the medium has repeatedly been refreshed [21], while in experiment II this has not been done so often (private communication), leading to lower concentrations of nutrients and other molecular factors in experiment II. During the whole course of osmotic stress application, an over-expression of the kinase inhibitor *p*27^*Kip*1^ together with an increased number of cells arrested in the G1 phase was observed, but no significant change in apoptosis rates after 3 days was reported.

(**EII.OIII**) Delarue et al. (2014) also considered the stress response for other cell lines (AB6, HT29, BC52, FHI) performing steps EII.OI and EII.OII for each cell line. These data will be used to validate our model despite less information concerning cell size and cycling times is available for these cell lines.

### Hypotheses for growth and death of tumor cells

As a first step we proposed a number of hypotheses for the growth dynamics common to experiments I and II.

(**H.I**) In both experiments a linear growth phase was observed after exposing the MCS to external stress. The growth of the cell population that is not constrained by either mechanically-induced growth inhibition, nutrient, oxygen or growth factor limitations is exponential [4]. We assumed that deviation of growth from an exponential indicates limitation of proliferation to a proliferating rim. This may have different reasons, for example necrosis that has been only reported for experiment I (EI.OIV), or of cells being quiescent. Both necrosis and quiescence can result from a lack of nutriments or other factors [6,24], that may indirectly be promoted by pressure, e.g. in case the compression of the cell layer squeezed between the capsule shell and the inner cell layers leads to the formation of an obstructive barrier for some nutriments (as glucose) to the cells located more deeply in the interior of the tumor. However, cell quiescence or cell death may also be a direct consequence of mechanical pressure, e.g. if cells subject to compression cannot advance in cell cycle for too long and then undergo apoptosis [6, 24]. We do not specify the origin the proliferating rim here, we take it into account through the definition of a thickness λ_*k*_(*k* = *I, II* is the experiment index). In Exp. I, λ_*I*_ distinguishes the necrotic cells from proliferating ones, in Exp.II, λ_*II*_ separates the quiescent from the proliferating ones. Necrotic cells as observed in experiment I can undergo lysis, in which they steadily lose a part of their fluid mass. The decrease of mass is limited to about 70% – 90% of the total initial mass of the cell [25,26].

(**H.II**) Cell growth rate may be declined or inhibited by pressure [8]. The authors of a recent study [12] hypothesized that the growth rate may be down-regulated if the cell volume is reduced as a consequence of pressure. We here test the hypothesis that growth rate is dependent on the volumetric strain (“true strain”, commonly used in case of large strains),

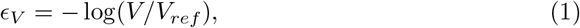

where *V* is the actual compressed volume and *V_ref_* is the volume of the cell in free suspension. The volumetric strain can be related with the pressure by integration of the relation *dp* = −*Kdϵ_V_. K* is the compression modulus of the cell and depends on the actual volume fraction of water, and the elastic response of the cytoskeleton structure and organelles. It may also be influenced by the permeability of the plasma membrane for water, the presence of caveolae [27], and active cellular responses. As such, the timescale at which *K* is measured is important.

In our simulations, we regarded *K* as the long timescale modulus of cell, as growth and divisions are slow processes. We studied constant and a volume-dependent compression moduli (the calculation of growth, volume and pressure for each cell in the model is explained in the Methods section Cell growth, mitosis, and lysis, Equation 8).

On the molecular level, volume reduction correlates with over expression of *p*27^*Kip*1^ which progressively decreases the proliferating potential. Other molecular players such as the transcriptional regulators YAP/TAZ were also reported to be mechano-sensitive [28]. In the scope of the present work, these reports suggest that quiescence, and perhaps also apoptosis, may be controlled by either pressure or cell volume. Experimental studies [29–32] mainly measured the growth rate of dry mass or size. These indicate that the growth rate *α* varies within the cell-cycle, yet a unique relationship is difficult to infer.

We propose as general form for growth rate *α* the Hill formula:

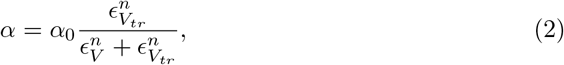

where *α*_0_ is the growth rate of the unconstrained cell, *ϵ_V_tr__* is a threshold value^1^, and *n* is an integer. The parameter *ϵ_V_tr__* is the value where the cells have lost 50% of their initial growth rate. Note that for *ϵ_V_tr__* → ∞ we retrieve a constant growth scenario, whereas increasing *n* from 1 to ∞ modifies the curve from a linear-like decrease to a sharp pressure threshold (see Figure 2A). The use of a Hill function thus makes a variety of growth scenarios possible. Hill formulas have been used in the past to simulate contact inhibition in epithelial tissue and tumors [17,33,34]. We discuss the generality of this approach in the Discussion section.

**Fig 2.**
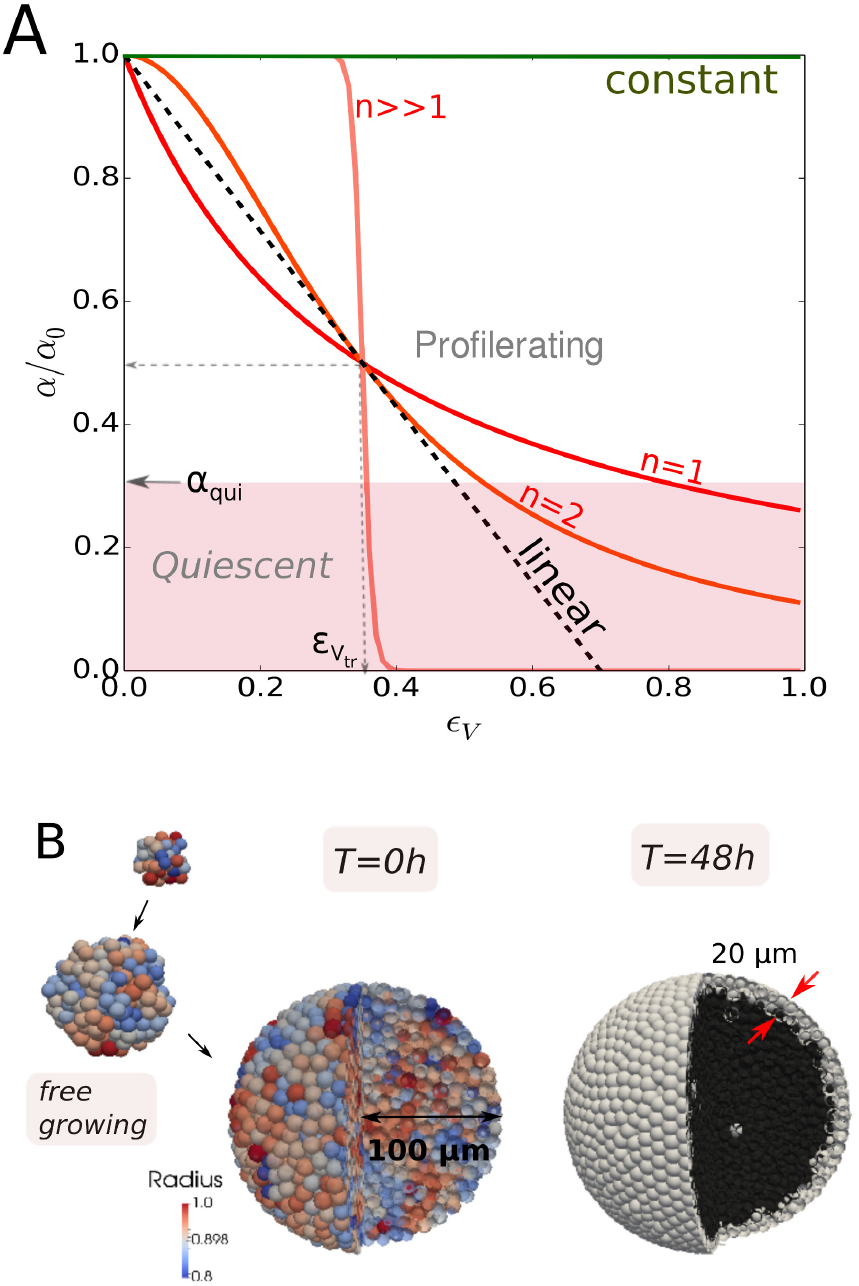
(A) Plot of Hill growth rate function as function of the volumetric strain *ϵ_V_* = *ϵ_V_*(*p*), for *n* = 1, 2 and a large value of *n*, and for a constant growth scenario (*ϵ_V_tr__* → ∞). Plot of a linear growth rate function with *ϵ_V_tr__* such that *α/α*_0_ = 1/2. Below the pink zone indicated by *α_qui_* cells become quiescent and growth stalls. (B) simulation snapshots of a CT26 spheroid during the initial free growth, just before confinement (coloring according to cell radius), and at 48h of confinement in capsule (coloring here indicates necrotic cells (dark) and viable cells (white)).

(**H.III**) It is generally accepted that cells who have passed the G1 checkpoint (also known as restriction point) are committed to divide, else they go into quiescence (G0). In our model we assume this checkpoint is situated after 1/4 of the total cell cycle time [35]. The transition criterion to the quiescence state can be defined as the one at which the growth rate “stalls”, i.e. *α*/*α*_0_ < *α_qui_* (see Figure 2A).

“Sizer versus Timer”: According to hypothesis H.II growth rate depends on the compression of the cells, hence the volume doubling time can locally vary and is larger than for uncompressed cells. Limiting cases would be that division occurred after volume doubling at a variable time [6] (“sizer”), or after a pre-defined time (“timer”) often mentioned in developmental biology [36]. We therefore also compared the effect of constant time vs. doubling of volume criterion in cell division on the cell population behavior. Also mentioned in H.II, the unconstrained growth rate *α*_0_ itself may vary during the cell cycle. To study the potential effect of these variations we performed comparative runs considering constant growth rate as well as exponential growth rate during the cell cycle (details in section Cell growth, mitosis, and lysis).

### Establishment of the Agent-Based Model and its parameterization

For the model development and parameterization we pursued a multi-step strategy sketched in Figure 3 (see also Table 1 and 2). The model parameters for the “model I” to mimic experiment I, 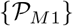, and “model II” to mimic experiment II, 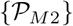, were step-wise calibrated from experiments I and II, and in each case first for growth in absence of external mechanical stress on the growing population, then in presence of stress. They can be categorized by separating between cell line-specific parameters 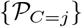, where *j* ∈ {*CT*26, *AB*6, *HT*29, *BC*52, *FHI*}, determines the cell line, and experiment-specific parameters 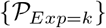 with *k* = *I, II* characterizing the experimental setting. The simulations were performed with a center-based model (CBM). As the model is parameterized by measurable physical and bio-kinetic parameters, parameter ranges could readily be determined within narrow limits (Table 2, [22]).

**Fig 3.**
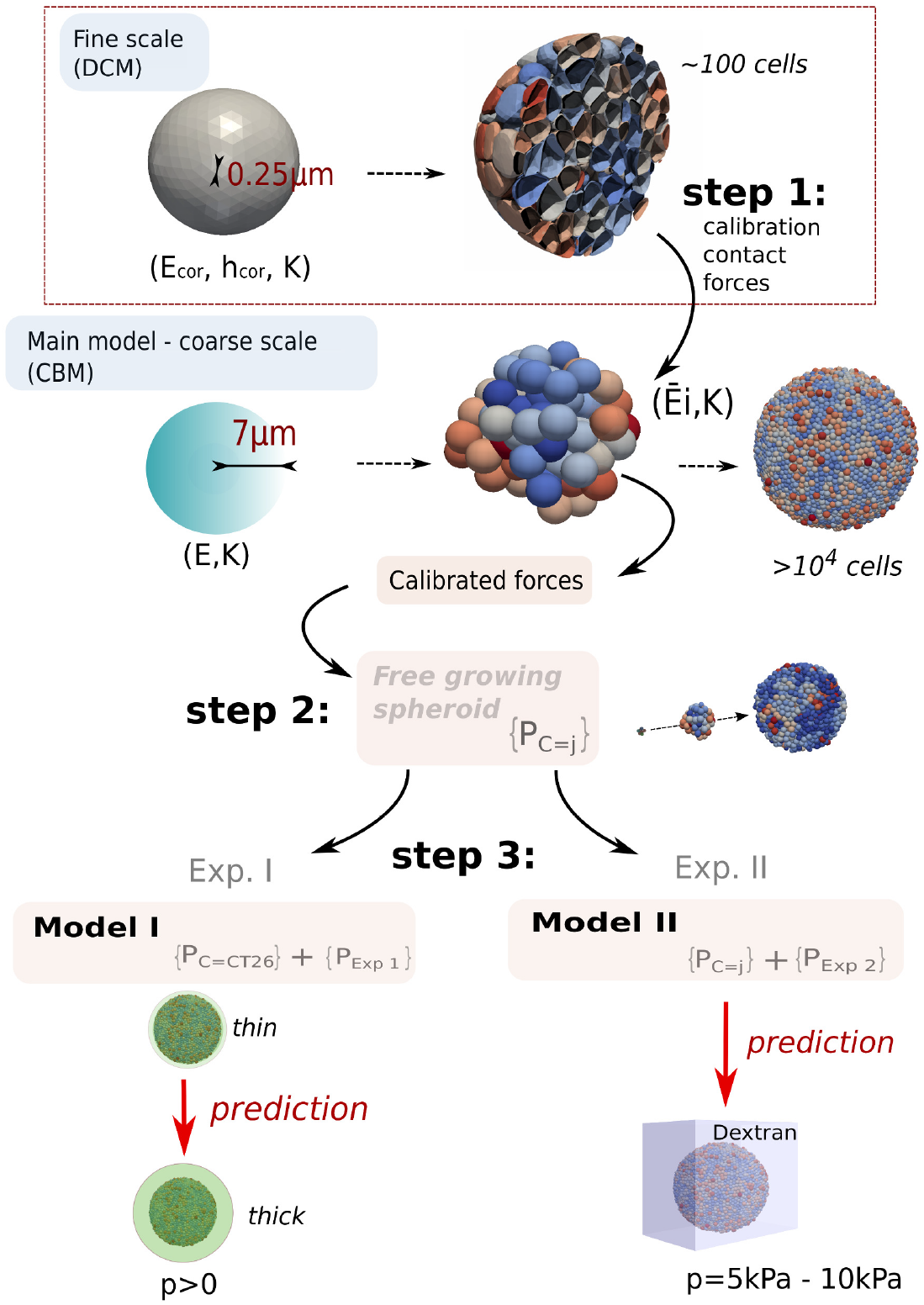
Model calibration overview. Simulations were performed with a center-based model (CBM). In step 1, the contact forces in CBM were calibrated from DCM simulations with parameters (*E_cor_, h_cor_, K*), yielding a variable effective contact stiffness 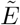 of the CBM depending on the compression level. In **step 2** the parameters 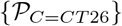 of the CBM for cell line CT26 were determined. Comparing simulations of the CBM with stress-free growth of multicellular CT26 spheroids in experiment I determines most parameters of 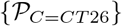 (Figure 1B, full black line). **step 3**: those cell-line parameters that are affected by the capsule, are specified by comparison with the data from experiment I in presence of the thin capsule. The set of experiment-specific parameters 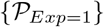 (Young modulus and thickness of the capsule) are given by the experimental setting. For the so specified complete set of parameters the simulation reproduces the experimental data I for the thin capsule (Figure 1B, dashed black line), and, after replacement of the capsule thickness, predicts the experimental data for the thick capsule (Figure 5B). For CT26 cells growing in experiment setting II the cell parameters remain unchanged 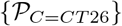. The deviation of the growth dynamics of stress-free growth from an exponential in experiment II (Figure 1B, full red line) is taken into account by an experiment-specific parameter, namely the proliferative rim. Without any further fit parameter, the model then predicts the correct growth dynamics subject to dextran-mediated stress (Figure 1B, dashed red line). In order to predict the stress-affected growth kinetics of the cell lines *j* = {*CT*26, *AB*6, *HT*29, *BC*52, *FHI*}, their cell cycle duration is modified to capture the stress-free growth analogously to that of CT26 cells in experimental setting II (Figure 1D-G, full red lines). After determining the parameters, the growth kinetics of these cell lines subject to stress could be predicted (Figure 1D-G, dashed red lines).

**Table 1.** Nominal physical parameter values for the DCM to calibrate the CBM.

| Parameter set | symbol | unit | value | ref |
| --- | --- | --- | --- | --- |
| Cortex Young's modulus | $E_{cor}$ | $Pa$ | 2400 | [37] |
| Cortex thickness | $h_{cor}$ | $\mu m$ | 0.1 | [37] |
| Cell compressibility | $K$ | $kPa$ | [2.5, 10] | CS, [12, 37–39] |

**Table 2.**
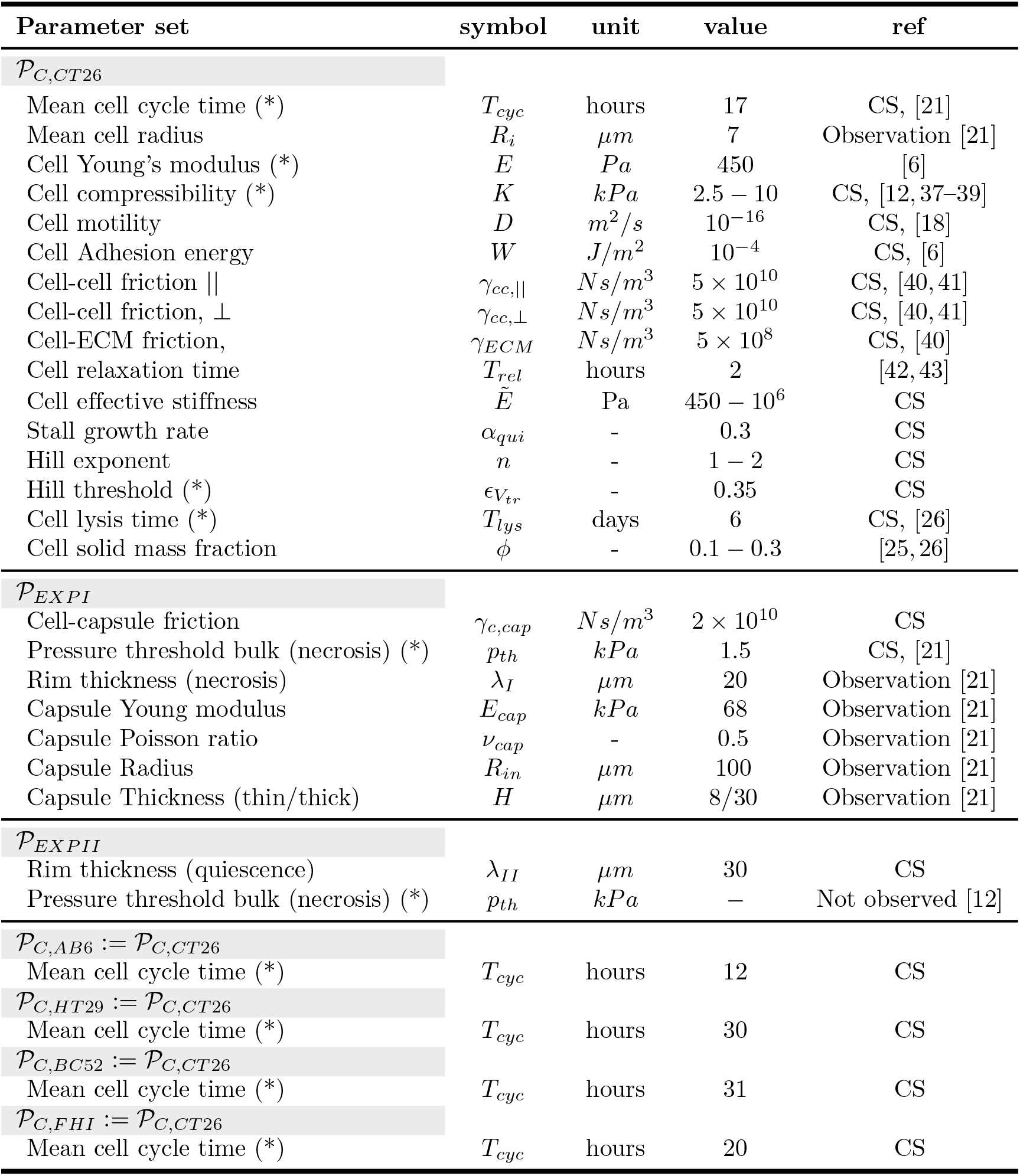
Reference physical parameter values for the model. CS indicates a model choice. If CS shows up with references next to it, the value was chosen from the parameter range in the references. A reference only means the value is fixed from literature. An (∼) denotes parameter variability meaning that the individual cell parameters are picked from a Gaussian distribution with ±10% on their mean value.

First 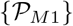 was identified in three steps (1)-(3) (Table 1).

(1) As the “standard” CBMs are inaccurate in case of high compression [20], the cell-cell interaction force in the CBM in this work was calibrated using computer simulations with a deformable cell model (DCM), resulting in an effective stiffness 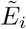 in the CBM at high compression, that increases with increasing compression, see Methods section Calibration of the CBM contact forces using DCM. 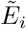 belongs to 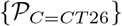 of the CBM. The DCM could not be directly used for the growth simulations, as it is computationally too expensive to run simulations up to the experimentally observed cell population sizes of ∼ 10^4^ cells. Next, the experimental information was taken into account (Figure 3).

(2) Comparing simulations of the CBM with the data from the stress-free growth control experiment of multicellular CT26 spheroids (MCS) in experiment I permits determining those parameters of 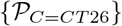 that were are unaffected by the presence of the elastic capsule (Table 2), see Methods section Model setup and parameter determination.

(3) Adding a thin elastic capsule specifies the set of experimental parameters 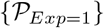 (Young modulus, Poisson ratio and thickness of the capsule etc.), and permits identifying those cell line specific parameters that respond on the presence of the capsule.

In experiment I these are the parameters characterizing cell cycle entrance and cell growth (2). Finally, model I is characterized by the conjunction of the cell-specific and the experiment-specific parameter sets 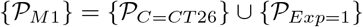.

Replacing the thin by a thick capsule in the simulations by changing the experimentally determined thickness parameter for the thin capsule in 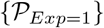 by that for the thick capsule leads to a predicted simulated growth dynamics that matches well with the one experimental data without any additional fit parameters (Figure 5B).

Experiment II has been performed with CT26, AB6, HT29, BC52, FHI cells. For CT26 cells, the cell-line specific parameter set remains the same in experiment II as in experiment I. Different from experiment I, stress-free growth in experiment II is not exponential but linear, reflecting different growth conditions that limit cell proliferating to a proliferating rim. This determines the proliferating rim size λ_II_ as the experimental parameter of set 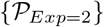 that summarizes the impact of growth medium under the conditions of experiment II in stress-free growth. In presence of dextran, 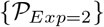 is expanded by only the measured pressure exerted by dextran, which as it is experimentally determined, is no fit parameter (λ_II_ remains unchanged). With the parameter set 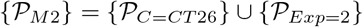, the simulation model predicts a growth dynamics that quantitatively agrees with the one experimentally found indicating that the growth response only depends on the exerted pressure, not on any other parameter (Figure 1B).

In a last step, the stress response of the other cell lines, *j* = {*AB*6, *HT*29, *BC*52, *FHI*} have been modeled for the experimental setting of experiment II, again in two steps (Figure 1D-G). The first step was to adjust the cell cycle time *T_cyc_* of the cell line to fit the stress-free growth leading to replacement of that one parameter in passing from 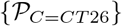 to 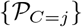, the second was predicting the growth subject to dextran-mediated stress without any parameter fitting i.e., using 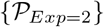 for the experimental parameters.

Summarizing, almost the entire parameter determination is done by adjusting the model parameters to experiment I for a thin capsule. After this step there is only one fit parameter for each cell line, summarizing the cell-line specific effect of growth conditions of experiment II for the stress-free growth (i.e., the control experiment). The step to simulate population growth subject to external stress, both in the thick capsule for CT26 as well as in experiment II with dextran for the cell lines CT26, AB6, HT29, BC52 and FHI is performed without parameter fitting.

### Model for experiment I with thin capsule

#### Calibration step

##### Growth without external stress

First, we simulated CT26 cells growing freely in the liquid suspension ((EI.OI), Figure 3) for the parameters, see Table 2). In this situation, CT26 cells grew approximately exponentially indicating absence of growth inhibition. For the simulation we needed to specify a subset of parameter set 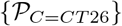, namely the division time *T_cyc_*, cell radius *R*, cell Young modulus *E* and cell compressibility *K*, characteristic lysis time *T_lys_*, the diffusion constant *D* of the cell as it specifies the micro-motility, the perpendicular and tangential cell-cell friction coefficients *γ_cc_*,∥ and *γ_cc_*,⊥, the cell-ECM (extra-cellular matrix) friction coefficient *γ_ECM_*, the cell relaxation time *T_rel_*, and the growth rate of the cell not subject to mechanical stress *α*_0_. For each of these parameters, either estimates from experiment I or literature estimates exist (see Methods section Model setup and parameter determination and Table 2).

For a constant cell cycle duration of *T_cyc_* = 17h (no inhibition), in the observation period −2*d* ≤ *t* ≤ 1d, we found a good mutual agreement between the model, the experimental growth curve, and an exponential, see Figure 1B. This determines the intrinsic cell cycle duration *T_cyc_* of a growing cell population subject to neither external mechanical stress nor nutriment limitation. (A movie (Video 1) of this simulation is provided in section Videos.)

##### Growth in presence of external stress

In the next step, we used the same model to mimic a growing multicellular spheroid in a thin capsule (*H* = 8 *μ*m). In the experiment after confluence, the growth curve crosses over into an approximately linear slope (*t* ≥ 1d in Figure 1B) at a measured pressure of *p_th_* ≈ 1.5k*Pa* (EI.OII) with a viable rim of size λ_*I*_ ≈ 20*μm* (see EI.OIV and H1) enclosing a necrotic zone. Necrosis indicates a lack of nutriments. It is possible that at that pressure, border cells may be so compressed that nutriment diffusion becomes inhibited.

As the experimental data needed to explicitly model the influence of nutrients is not available and would require knowledge on many parameters (see [24]), we do not model nutriment-dependency explicitly but directly implement the experimental observation that the cells further inside the capsule than at distance λ_*I*_ die at pressure *p* = *p_th_* (observation EI.OII and Figure 4C), see section Model setup and parameter determination for more details.

**Fig 4.**
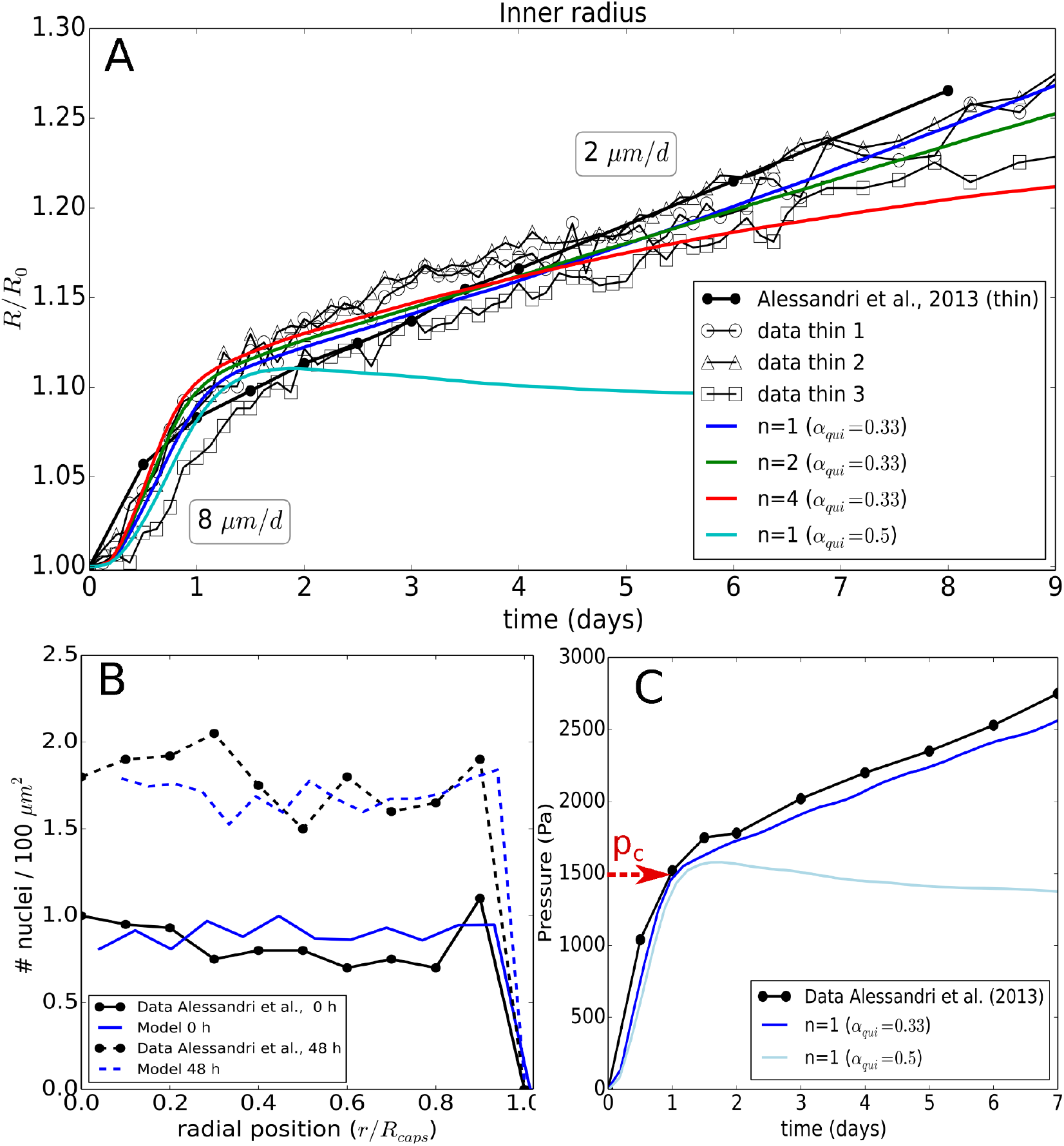
(A) Time evolution of the radius of the thin capsule for the experimental data and the simulations using Model I showing the effect of a parameter variation for n with *α_qui_* = 0.33, and *n* =1 with *α_qui_* = 0.5. (B) Simulation and experimental values of the radial cell density in the spheroid at *T* = 0*h*, and *T* = 48*h* for the optimal parameters. (C) Pressure curves indicating the pressure at the transition point from free spheroid growth to spheroid growth against the thin capsule in ref. [21] and the simulation.

In our first attempts all cells in the viable rim were assumed to proliferate with a constant rate *α*_0_. This assumption led to a too high spheroid growth speed, hence could not explain the growth kinetics in presence of the capsule (see section Model setup and parameter determination, Figure 10A), expressing that λ_*I*_ does not determine the growth speed, but only the size of the viable rim.

The constant growth speed for *t* > 2*d*, despite increasing pressure experienced with increasing size of the MCS, indicates the proliferating rim to be of constant size. This was confirmed by visual observation of the spheroids (personal communication). This argues against an increasing limitation of nutriments with tumor size in the linear growth regime, and in favor of a direct impact of pressure on cell cycle progression.

In our model this was taken into account by replacing the constant growth rate *α*_0_ by a compression-dependent growth rate *α*(*ϵ_V_*) Equation 2 expressing, that cells can enter G0 if the relative growth rate *α/α*_0_ falls below a threshold *α_qui_* between division and restriction point, see H.III and Figure 2). In our model, cells divide after their volume have doubled. Consequently, a cell subject to compressive stress has a longer cell cycle duration than an isolated cell.

With this model we found a very good agreement between experimental data and simulation results for *ϵ_V_tr__* ≈ 0.35, *n* ∈ [1, 2] and *α_qui_* ≤ 0.33 (Figure 4A, Figure 4B)). Values of *n* ∈ [1, 2] do hardly discriminate. Choosing *n* ≥ 4 results in a faster growth in the beginning as here *ϵ < ϵ_V_tr__*, and an experimentally not observed flattening of the residual growth resulting from the sharp decrease of *α* for *ϵ_V_* > *ϵ_V_cr__. n* → ∞ leads to a plateau. Increasing *α_qui_* to 0.5 results in a significant growth stall as cells then already enter quiescence at higher growth rates (Figure 4A). Increasing *ϵ_V_tr__* results in a faster capsule dilatation over the whole period as then the growth rate decreases only above a larger pressure (noticing that *dϵ_V_/dp* > 0). We selected *ϵ_V_tr__* ≈ 0.35 as best fit. The effect of *ϵ_V_tr__* is shown in the thick capsule experiment (section Validation of model for experiment I with thick capsule data, Figure 5A). The Hill function parameters complete parameter set {*P*_*C=CT*26_} (Table 2).

**Fig 5.**
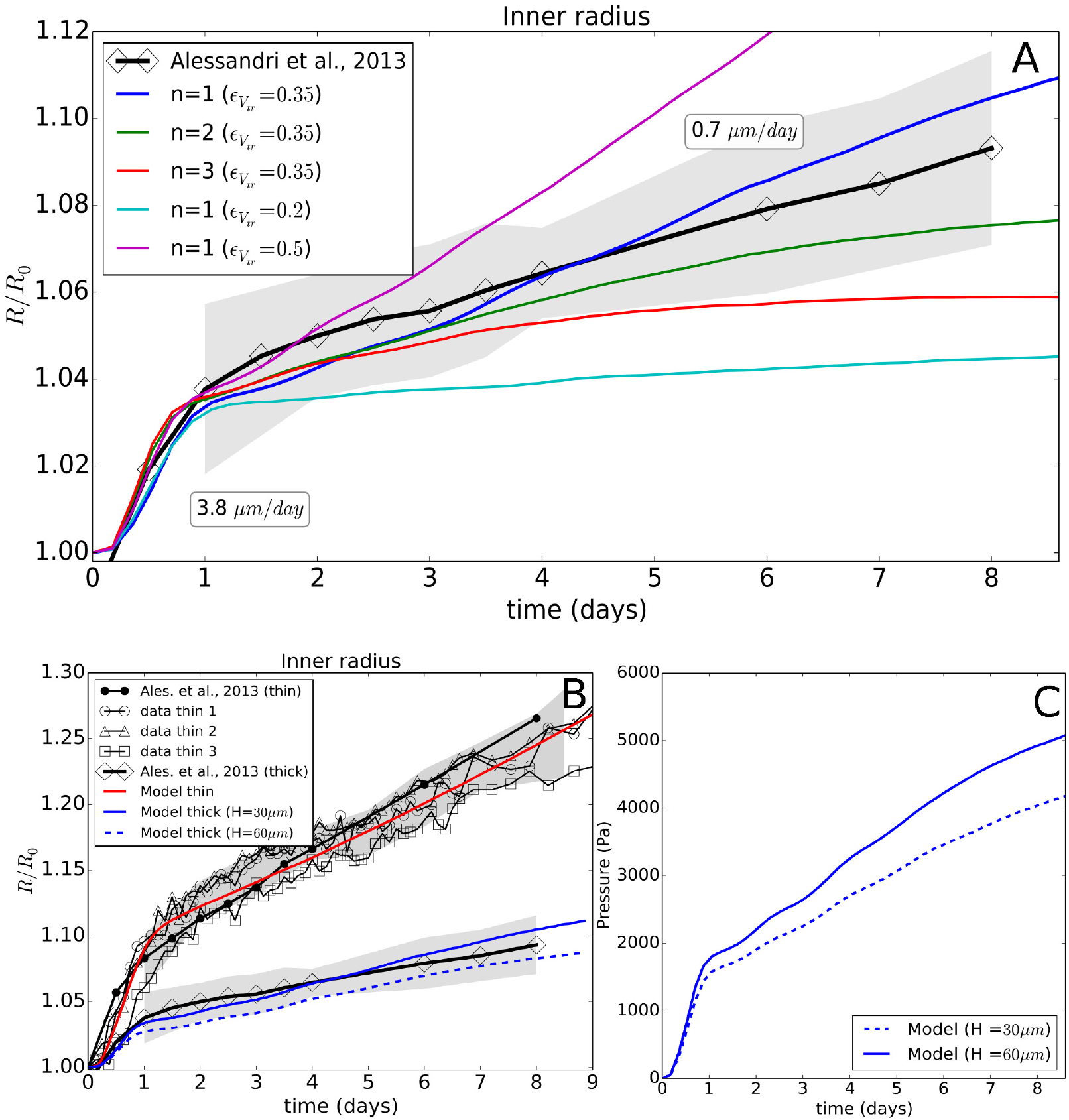
(top) (A) Time evolution of the thick capsule radius (*H* = 30 *μ*m), shown for the experimental data and the simulation with Model I, indicating the effect of the parameter *n* and *ϵ_V_tr__*. As the number of data sets on the thick capsule did not suffice to estimate the experimental error, the errors on the thick capsule data (gray zone) were estimated from the spreading on the thin capsule data, by determining the minimum - maximum intervals for the thin capsule data. These were then rescaled by the ratio of thin - thick capsule dilatations and shifted on to the thick capsule curve. (B) Global view of experiment I and II and respective model runs, including a model prediction for a capsule wall thickness *H* = 60 *μ*m. (C) Simulated evolution of the average pressure in a capsule with *H* = 30 *μ*m and *H* = 60 *μ*m.

We verified that the replacement of *α*_0_ by *α*(*ϵ_V_*) did not result in a disagreement between model simulation and experimental data for stress-free growth (black full line in Figure 1B) indicating that no critical pressure builds up for MCS growth in liquid suspension in absence of the capsule during the experimental observation time period.

We have also tested the hypotheses whether cells either have a growth rate *α*, constant during the cycle, or an exponential increase (see Methods section Cell growth, mitosis, and lysis), yet we did not find any significant differences for the spheroid growth, indicating robustness of the results against such variations.

As an alternative mechanism to cell division after volume doubling we also tested the assumption that a cell rather divide after a fixed cell cycle time (“timer”). This resulted in smaller daughter cell volumes if the mother cell experienced compressive stress during growth, and as a consequence in a too large nuclei density at 48h (see Figure 10C).

Concluding, using Model I a good agreement with data could be obtained whereby the main underlying assumption is that the cell growth rate and thereby the duration of the cell cycle is controlled by the cells’ degree of volumetric compression. (A movie (Video 2) of this simulation is provided in the Supplementary Material, section Videos.)

### Validation of model for experiment I with thick capsule data

In the first validation step, we considered the thick capsule experiment (*H* = 30 *μ*m). A thicker capsule provides a stronger resistance against the spheroid expansion. In simulations with model I and the parameter set (*n* ∈ [1, 2], *ϵ_V_tr__* = 0.35, *α_qui_* = 0.3) that was able to explain the MCS growth against a thin capsule, we obtained a good agreement also for the thick capsule data without any additional fit parameter (Figure 5A).

For higher or lower values for the volumetric strain threshold *ϵ_V_tr__*, respectively, an overestimation or underestimation for the residual growth would be observed consistently with the thin-capsule data. Values *n* ≥ 2 resulted in a clear deviation the end of the observation period and were hence rejected.

In the work of Alessandri et al., additional experiments were performed using thick capsules with a larger sizes (*R*_0_ ∼ 400 *μ*m) and thicker walls yet with the same aspect ratio *H/R*_0_ ∼ 0.25. The experiments show that the presence of a capsule did not affect the free growth of the MCS. The growth dynamics after confluence for the large thick capsule could not be uniquely determined as the duration of this phase was too small. For this reason we here did not simulate this case (see Figure 12F). Yet, to permit further validation of the model we also depict simulations for a capsule with thickness *H* = 60 *μ*m. This run predicts a slightly lower dilatation rate (Figure 5G) yet the pressure increase per day in the capsule (Figure 5C) is comparable with the 30 *μ*m case, about 250 Pa/day.

### Validation of model for experiment II: same cell lines as for experiment I

#### Model II

We challenged the model calibrated for experiment I by studying whether it would be able to predict the observed growth of CT26 multicellular spheroids subject to osmotic stress (Experiment II, [12]). The concentration of dextran regulates the applied pressure. The growth rate at *p* = 5 kPa here is also significantly lower than those in control spheroids (freely growing in iso-osmotic conditions). Surprisingly however, the control spheroids in experiment II grow slower than in Experiment I, revealing an overall linear but not exponential growth kinetics. Since the cell line is identical, we associate this difference to varying culturing conditions (e.g. less frequent change of medium).

##### Growth without external stress

To take the different culture conditions into account within our simulations, we first simulated again the free growing spheroid. Linear growth is characteristic for a proliferative rim of constant size, with the size and spatial distribution of proliferating cells in the rim determining the speed of spheroid expansion [24, 44]. Following the same reasoning as for experiment I, we impose a proliferating rim of size λ_*II*_ measured from the edge of the spheroids inwards to capture the linear growth. Here, the edge of the spheroid is computed as the average of the radial positions of the most outer cells plus one mean cell radius (see Figure 6A). We found that for *λ_II_* = 30 *μ*m with cells adopting the same parameter set as in Experiment I, Model I (*n* =1, *ϵ_V_tr__* = 0.35, *α_qui_* = 0.3), matches well with the data for freely growing spheroids (Figure 7A). As in experiment II no increase in cell death, neither by apoptosis nor by necrosis has been reported, cells outside of the proliferating rim are assumed to rapidly enter a quiescent state without undergoing necrosis i.e., they do not shrink. This is referred to as Model II. Notice that λ is the only parameter value by which Model II differs from Model I, reflecting the response on the growth conditions (therefore attributed to the parameter set 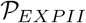).

**Fig 6.**
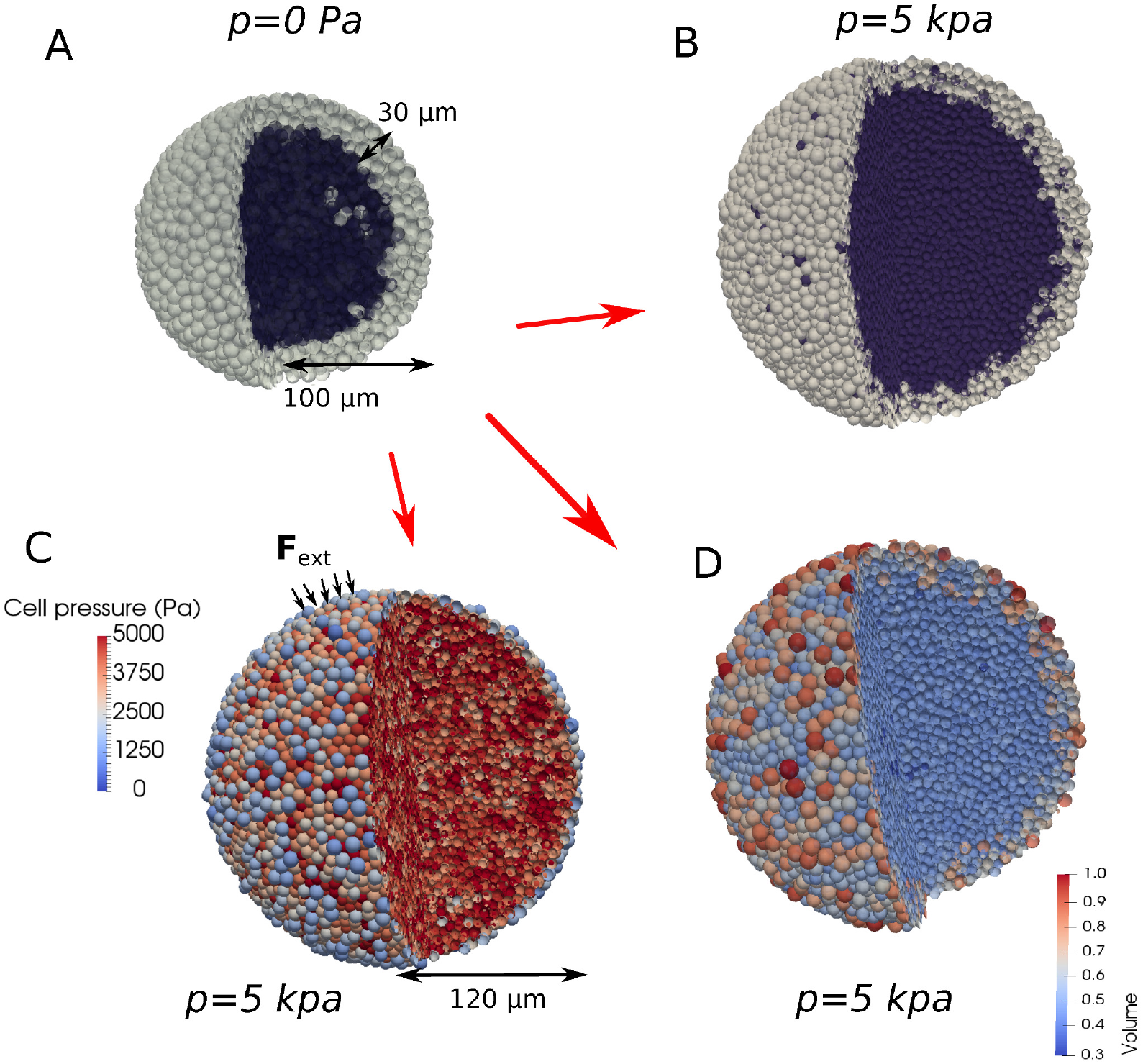
(A) Simulation snapshot at the beginning of a free growing CT26 spheroid (*R* = 100*μ*m), indicating quiescent (dark) and proliferating cells (light). (B-D) Simulation snapshots of growing CT26 spheroids at *R* = 120 *μ*m during dextran application (*p* = 5kPa), indicating quiescent and proliferating cells (B), individual cell pressure (C), and volume for the cells (D).

##### Growth in presence of external stress

The same parameter values are kept for the growth simulations in the presence of dextran. In another work by Delarue et al. (2014) [38], slight cell elongations were reported towards the tumor center. We neglected here this effect to test whether the experimentally observed response of a growing tumor subject to osmotic stress can already be captured with the model originally developed for the capsule, with the only difference being an adaptation for the free growth conditions.

In accordance with the known pressure-exerting effect of dextran, we apply an external force only to a small boundary of outer cells, directed towards the center of the spheroid, mimicking the osmotic effects which induce depletion-induced adhesion and an increase of the contact area between the cells [45]. The magnitude of the applied force on every outer cell reads:

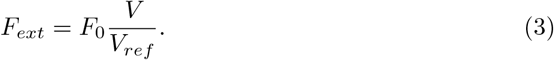

The magnitude *F*_0_ (fixed parameter) is chosen such that the experimentally observed average cell pressure 〈*p*〉 is in the simulation maintained in the bulk of the spheroid during growth. The volume-scaling factor is needed to minimize pressure variations as much as possible. As there is no confining volume of the MCS, we use a *local* calibration approach to compute the contact forces in the agent-based model, see section “Local” calibration approach, needed for experiment II.

Remarkably, the slope of the growth curve obtained from a simulation with the model without any further adjustment matches very well with the data (Figure 7A and Figure 1B). This indicates that the response of the CT26 cells on compressive stress is robust and reproducible even if the cells are subject to different environmental conditions. Moreover, the surprisingly good agreement between model prediction and experimental observation suggests that the slight cell elongations observed in [38] might not be a fundamental determinant in the overall response of a growing tumor to external mechanical stress by osmosis. The major contribution to the stress response may be controlled by the proliferating cells that are mainly located close to the border. As proliferating cells, which are on average larger than resting cells, are mainly localized at the border, the nuclei-nuclei distance is larger close to the border of the spheroid than inside (see Figure 6D), consistent with reported experimental observations in [12] and in freely growing spheroids [44].

**Fig 7.**
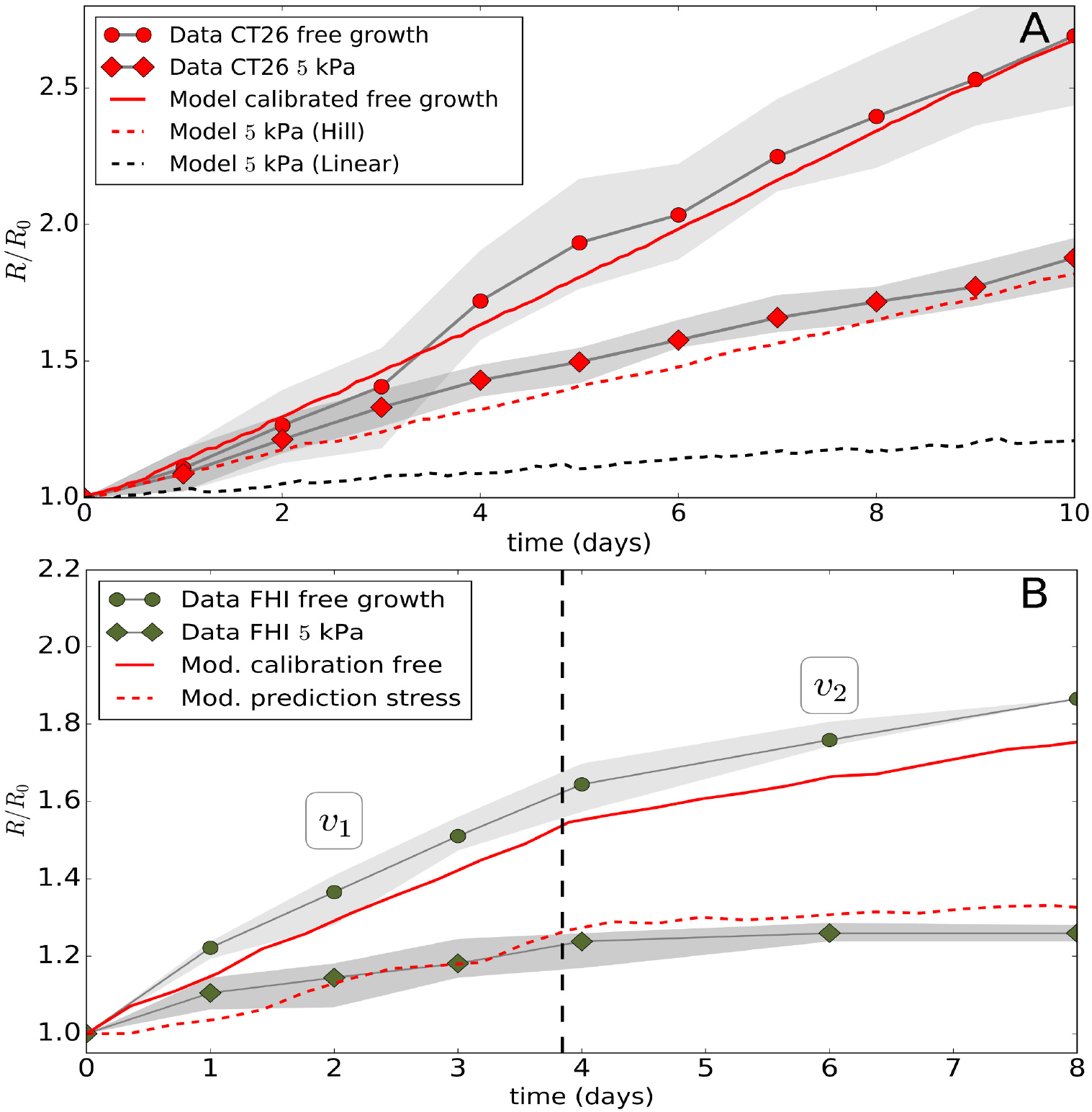
(A-B) Detail of the time evolution of radius of the CT26 and FHI spheroid relative to its initial state. Data from [12] shown for free growth and at *p* = 5 kPa. Runs with Model II are for free growth and for *p* = 5 kPa. In the CT26 cell line an additional model run is shown assuming a linear cell cycle progression function. In the FHI cell line the vertical line indicates the presumed changes in experimental conditions for free growth over time resulting in a lower surface growth (*v*_1_ → *v*_2_). The gray zones in the plots indicate the min-max values of the data.

Within our model we find that i) the pressure distribution in the bulk cells is quite homogeneous, and ii) the pressure is locally lower for the most outer cells because some of these cells are experiencing less contact forces from their neighbors (see Figure 6C).

In simulation runs testing parameter sensitivity of the growth kinetics in Experiment II we found for growth parameters *α_qui_* > 0.33, *ϵ_V_tr__* < 0.2 or *n* > 2 a significant underestimation of grow (too many cells go into quiescence), in agreement with our simulations for Experiment I.

### Validation of model for experiment II: other cell lines

In order further challenge our model, we also simulated the dextran experiments performed with other cell lines, i.e. AB6 (mouse sarcoma), BC52 (human breast cancer), FHI (Mouse Schwann) all at *p* = 5 kPa, and the cell line HT29 (human colon carcinoma) at *p* = 10 kPa. Since these experiments were less documented, our assumptions are that i) in the simulations the experimental conditions are a priori the same, but ii) cell cycling times are different. These doubling times were estimated by calibration of the growth curves without external stress before predicting the growth curves in presence of external stress without any additional fit parameter following the same strategy as for experiment II above for the CT26 cell line. Doing so, we found that the long-term growth speed was again surprisingly well predicted by the model for all three cell lines. Only transients partially deviate from experimental curves (Figure 1D-G, Figure 7A-B).

We here adjusted the cell cycle duration *T_cyc_* to capture the growth kinetics of the MCS in absence of externally exerted mechanical stress but we could also have modified, for example, the thickness of the proliferating rim λ_*II*_, as the expansion speed *v_f_* of the freely growing MCS is *v_f_* ∝ λ_*II*_/*T_cyc_* [46], so that changing λ_*II*_ has the same effect as the opposite change in *T_cyc_*. We emphasize in this context that λ_*II*_ does not determine the growth speed *v_S_* under dextran-induced stress, as *v_S_* ≪ *v_f_*. Thus, our prediction is not dictated by parameter λ_*II*_.

For AB6 (Figure 1E), we found a doubling time of 13*h* to make the simulated free growth case matching well with the experiment (comparing slopes over period of ∼ 9*d*; full red line in Figure 1E). We however, did not have any additional information concerning cell size and doubling time on this cell line. Applying the pressure of 5*kPa* in the simulations, one still sees that the simulation agree quite well with the experiment (Figure 1E, dashed red line).

For HT29 (Figure 1G), a pressure of 10 kPa was applied in the experiment, and hence this puts an extra challenge as the growth model is tested for larger compression. In the simulations, we now had to double the applied forces in the most outer cells to reach the same average pressure. The calibrated doubling time of HT29 for growth in absence of dextran was found to be 46*h*, in agreement with values in reported in [47] (full red line in Figure 1G). The cell size is comparable to that of CT26 [12]. The simulation results in presence of dextran indicates a significant differences in the beginning of the experiment, yet overall the growth slope matches quite well with the data (Figure 1G, red dashed line).

Finally, for BC52 (Figure 1D) and FHI (Figure 1F and 7B), the experimental results show a more complex behavior, as there seem to be two regimes in the growth. In the case of BC52 the spheroid first grows with *v*_1_ ∼ 0.41 *μ*m/h for the first 9d, then in the subsequent period the growth slows down to *v*_2_ ∼ 0.29 *μ*m/h (see Figure 1D). We attributed this to a change in growth conditions in the experiment. The model a-priori does take the cross-over effect into account, but we still can test it by imposing ad-hoc changes of experimental conditions after a period of 9*d*. To so so, we assumed in the simulations for the dextran-free growth that the thickness proliferating rim has decreased during the cross-over by λ_*II*_ → λ_*II*_ × *v*_2_/*v*_1_ ≈ 0.7λ_*II*_, which resulted in an overall good calibration curve (full red curve in the Figure 7B). The same procedure was applied to the FHI cells, with here the factor *v*_2_/*v*_1_ ≈ 0.35 for the simulation in absence of dextran (see full red line in Figure 7B). The corresponding simulations in presence of dextran for BC52 (Figure 1F, dashed red line) and FHI (Figure 7B, dashed red line) then shows that the model is again able to predict the experimentally observed slopes in both regimes reasonably well.

Hence, we conclude that this model is able to predict the effect of mechanical stress on the expansion speed of the MCS in the elastic capsule experiment (experiment I) and the dextran experiment (experiment II) after calibration of the model parameters with experimental growth data in absence of capsule and dextran i.e., with experimental growth kinetic data in absence of externally exerted mechanical stress.

### Robustness of the proposed cell cycle progression function

In our model we had proposed that the cell growth rate decreases according to a general Hill function (Equation 2). From the capsule simulations, we observed that neither a constant growth scenario (*ϵ_V_tr__* → ∞) nor a sharp threshold (*n* → ∞) could explain the data. However, in order to justify the choice of the Hill functional shape as compared to a simpler functions, we have performed comparative simulations with a linear progression function. This function has the same boundary value *α* = *α*_0_ at *ϵ_V_* = 0, and *α* = 0.5 × *α*_0_ at *ϵ_V_* = 0.35, but has a steeper decrease further on (dashed line in Figure 2). We found that with this function the experimental data for small and large capsule thickness could still be reproduced with a fair agreement (see Figure 12E, “Linear I” in the Appendix). However using the same function, we could subsequently not match the data of Experiment II, for the CT26 cell lines as well as for the other cell lines. In that case the simulations systematically underestimated the growth (see Figure 7A, black line) indicating the tail of the Hill function is important as it controls the still non-negligible contribution to growth at high strains occurring in the dextran experiment. On the other hand, a linear function (boundary value *α* = *α*_0_ at *ϵ_V_* = 0) calibrated such that the CT26 dextran experiment could be reproduced, resulted in an overestimation of growth in the capsule experiment (see Figure 12E, “Linear II” in the Appendix). Concluding, a sufficiently long “tail” in the diagram *α* versus *ϵ_V_* seems to be necessary to explain the residual growth of the cells. This points towards an nonlinear response of inhibition of growth of the cells upon compression, and further shows that the choice of a nonlinear progression function is necessary so that a Hill growth function, despite it looks complex, seems the most simple one that is able to explain simultaneously growth of MCS subject to externally applied stress in both experiment types.

## Discussion

By establishing a quantitative model of growing multicellular spheroids (MCS) subject to compressive stress calibrated with data on growth in an elastic capsule we were able to demonstrate that the stress response of a growing tumor is quantitatively robust and reproducible even if cells grow under different conditions and if the pressure is exerted by different experimental methods. Given the enormous complexity of intracellular processes involved in the control of MCS growth this is fascinating as it might open the possibility that largely separated robust functional modules may be identified and studied in separation without the need to analyze all interactions of the components of one module with the components of other modules, and without incorporating all interactions at the molecular level. In particular, we first developed a model to study CT26 cells grown in an elastic thin and thick capsule, and then modified this model in a minimal way by taking into account the remarkably different growth behavior of freely growing tumor spheroids (i.e. not subject to compressive stress) to simulate the tumor growth response of CT26 and other cell lines in a dextran solution. We show that the mechanical stress response is quantitatively the same despite significantly different culture and protocol conditions. Without the model, it would have been very difficult to identify this equivalence. The key results of our analysis are:

(**R.I**) With increasing compression the cell growth rate decreases. This relation could be well captured by a Hill function for the growth rate *α* that depends on the volumetric strain (Equation 2), and a transition into quiescence if the growth rate dropped below a threshold value. A sharp volume or pressure threshold below which no cell cycle entrance would occur anymore, is not compatible with the data. Together with the strain hardening assumption of cells during compression, this overall points to a nonlinear increasing growth resistance of the cells upon mechanical stress.

(**R.II**) Cells divide when their dry mass has doubled during the cycle. A “timer” as a decision mechanism for dividing could not explain the data.

A particular point of concern in many studies of spheroids is the appearance of cell death. Our work is based on the observations of Alessandri et al. (2013), who observed necrosis (CT26 cells, using FM4-64) in capsule confined cells, while their free growing spheroids exhibited the normal exponential growth for *R* < 150 *μ*m. Helmlinger et al. (1996) [8] observed a decrease in apoptotic (LS174T cells, using TUNEL) events during compression, and reported little necrosis (not quantified) for spheroids with *R* < 150 *μ*m. They concluded that the haltered growth of the spheroids is mainly due to the increasing compressed state, which can be partially confirmed by our simulations. In the work of Delarue et al. (2014) [12], no increase of apoptosis (HT29 cells, using cleaved-caspase 3) was observed after 3 days for spheroids with *R* ∼ 100 *μ*m. Contrary, earlier Montel et al. (2012) [11] did report increased apoptosis using cleaved-caspase 3 for CT26 cells, while Cheng et al. (2009) [9] did observe an increase of necrosis (67NR cells, using propidium iodide) even in very small spheroids *R* ∼ 50 *μ*m, yet mainly for the interior cells. At the periphery, cells were still dividing. Whether necrosis and apoptosis occurs may well be dependent on the cell type and experiment, but overall it seems that the peripheral cells are unaffected.

Our modeling strategy is based on *in silico* experiments i.e., abstracted experiments on the computer, where each individual cell was represented as modeling unit with those properties, actions and interactions that were considered as necessary to quantitatively explain the cellular growth response on mechanical compression. The implementation of cell-cell and cell-environment interaction directly accounts for physical laws with (in principle) measurable physical parameters that permit straightforward limitation of parameter ranges to those physiologically relevant. This made it possible for us to largely confine the parameter values to published or directly observed relatively narrow ranges, and introduce free fit parameters only for the cell cycle progression. A particular challenge was to construct an individual agent-based model that permits stable and robust simulations up to several tens of thousands cells under high compression. Under these conditions cell displacements may have to be minimal, which rules out models operating on lattices unless the lattice size would be chosen a very small fraction of the cell diameter (in which case they would lose their computational advantage). Thus, the requirements of constraining the parameters, and providing realistic simulation trajectories in time favored models operating in lattice-free space implementing a dynamics simulated by equations of motion (as opposed to a Monte Carlo dynamics, which under some condition mimics a master equation). The prototype of lattice free models are center-based models that calculate the forces between cells as forces between cell centers. However, as mentioned above and explained in more detail elsewhere [20] this model type has significant problems in dealing with cell populations under large compressive stress i.e., with exactly the situation we are faced with in this work. To solve this issue, we developed a deformable cell model, which represents each individual cell in much greater detail as in center-based models but at the expense of much longer simulation times. As simulations with that model up to several thousands of cells were not feasible, we performed simulations with this model of characteristic MCS configurations under large compressive stress and used the results to establish a new interaction force model within center-based models that permit to mimic large cell populations under large compression.

Finally, we mention that despite their limit on cell numbers, simulations with DCM can give valuable information on micro mechanics. In our study, we found that stiffer cells in a scaled capsule model more likely could cause a gradient in cell pressure from the border to the center of the spheroid than soft cells (see Appendix, Cell deformation and pressure distribution during in a compressed spheroid in DCM). These potential effects are difficult to investigate with center-based models and prove the necessity of further development of high resolution models, and perhaps running them on high performance computers.

## Mathematical methods for the agent-based models

This section summarizes the most important model assumptions and components, and then explains how model parameters were calibrated. More details about the mathematical formulations, can be found in the Appendix, section Appendix.

We start from a standard center-based model in which cells are represented by spheres. However, this model needs to be extended by calibration with a model that can deal with high compression, the “deformable cell model”, in order to obtain realistic results for the envisaged in vitro multi-cellular systems (section Calibration of the CBM contact forces using DCM).

### Center-based model (CBM)

In CBMs cells are approximated as simple geometrical objects capable of active migration, growth and division, and interaction with other cells or a medium [48]. In CBMs the precise cell shape is not explicitly modeled but only captured in a statistical sense. Here, the cells are represented by homogeneous isotropic elastic, adhesive spheres.

#### Equation of motion for the cells

The center of mass position of each cell *i* is obtained from a Langevin equation of motion, which summarizes all forces on that cell including a force term mimicking its micro-motility:

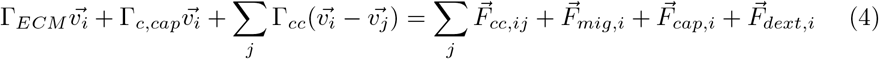

The lhs. describes cell-matrix friction, cell-capsule friction and cell-cell friction, respectively. Accordingly, Γ_*ECM*_, Γ*_c,cap_*, and Γ_*cc*_ denote the friction tensors for cell-ECM, cell-capsule, and cell-cell friction. The first term on the rhs. of the equation of motion represents the cell-cell repulsive and adhesive forces 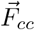, the 2nd term is an active force term 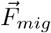, mimicking the cell micro-motility. 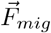 is mimicked by a Brownian motion term with zero mean value and uncorrelated in time (see Appendix). The existence of the 3rd and 4th term depends on the growth condition. In presence of an elastic capsule as in experiment I, the 3rd term denotes the interaction force experienced by the cell from the capsule 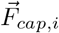 for those cells *i* that are in physical contact with the capsule. As cells cannot adhere to the capsule, 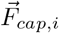 is purely repulsive. In absence of a capsule this term is dropped, 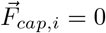. Analogously, in presence of dextran, 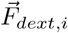 denotes the body force induced by dextran on the outermost cells i. In absence of dextran, 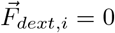. Due to high friction of the cells with their environment, inertia is neglected [49]. Based on the observation that ECM is produced by the cells (visualized by fibronectin staining [21]), which forms a substrate for the cells to actively migrate before confluence is reached, the first term on the lhs and the 2nd on the rhs express interactions with ECM. As the ECM network form fibronectin indicates a mesh size of the order of the cell size, and the ECM stiffness is usually much higher than that of cells [73], we assume momentum transfer to the ECM by the ECM friction and active micro-motility term but we do not model the ECM explicitely. When confluence is reached, the expansion of the spheroid originates from the volume increase of the cells against the mechanical resistance of the capsule or the osmotic forces, while the active micromotility force become negligible. This is further confirmed by simulations performing parameter variations in the micromotility forces which do not significantly influence the results (see section Model parameter and algorithm sensitivity).

#### Adhesive and repulsive forces

Interphase cells are approximated by homogeneous, isotropic, elastic and adhesive spheres which split into two adherent cells during mitosis. Under conditions met in this paper [40,48], the total cell to cell interaction force can be approximated by the sum of a repulsive and an adhesive force:

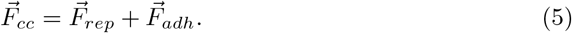

The repulsive Hertz contact force reads:

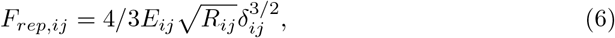

in which *E_ij_* and *R_ij_* are defined as

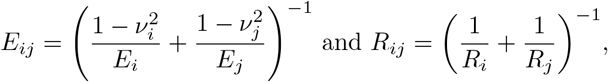

with *E_i_* and *E_j_* being the cell Young’s moduli, *ν_i_* and *ν_j_* the Poisson numbers and *R_i_* and *R_j_* the radii of the cells *i* and *j*, respectively. *δ_ij_* = *R_j_* + *R_i_* – *d_ij_* denotes the overlap of the two undeformed spheres, whereby 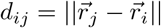 is the distance of the centers of cells *i* and *j* (see Figure 11A).

The original Hertz contact model does not take into account volume compression under large pressure by many surrounding cells. To account for multi-body interactions while using the classical Hertz model, we replace the Young moduli *E_i_* by an “apparent” contact stiffness 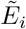 that increases as function of the cell density (Equation 14), see section Calibration of the CBM contact forces using DCM. The modification of the Hertz model is calibrated with a Deformable Cell Model (DCM) that represents cell shape explicitly.

The adhesive force term between cells can be estimated as proportional to the contact area and the energy of the adhesive contact *W* [50]:

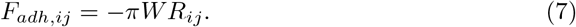

#### Cell volume and compressibility

In our model, cells are compressible meaning that cell volume is related to pressure by

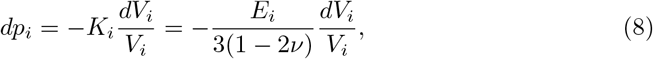

in case the cells’ properties are largely controlled by the elastic properties of its cytoskeleton and other cytoplasmic constituents. *K_i_* is the bulk modulus of the cell. The observed volume change in general depends on the speed of compression. For slow compression, water can be squeezed out of cells (and tissues), while for fast compression, water would yield incompressible resistance. In case *K_i_* = *K*_0,*i*_ is a constant, integration of the above equation yields the cell volume *V_i_* as a function of the pressure on cell i, *ϵ_V,i_* = (*p_i_* – *p*_0_)/*K*_0,*i*_ with *p*(*V_ref_*) = *p*_0_. Here, *ϵ_V,i_* = −log(*V_i_/V_ref,i_*) is the logarithmic strain permitting to capture large strains and 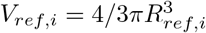 is the uncompressed cell volume the cell would have in isolation, with *R_ref,i_* being considered as constant for a quiescent cell. For small deviations *V* ≈ *V_ref_* the known relation *ϵ_V_* = log(*V*/*V_ref_*) ≈ (*V* – *V_ref_*)/*V_ref_* is recovered.

Several authors have reported strain hardening effects leading to an increased elastic modulus upon mechanical stress [51–53]. Stiffening of the cells can occur as the cytoskeleton gets denser [54]. In case of strain hardening, *K* increases with decreasing volume. We mimicked this by [54]:

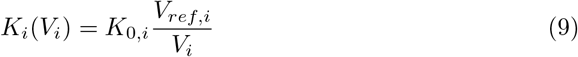

with *K*_0,*i*_ the compression modus of cell *i* in absence of stress. In this case, *ϵ_V,i_* = log((*p_i_* – *p*_0_)/*K*_0,*i*_ + 1). The quantity of interest is the volume response on a pressure change *p_i_* – *p*_0_, whereby throughout this paper we set *p_i_* ≡ *p_i_* – *p*_0_.

Now we assume that as a consequence of internal friction and by remodeling of the cytoskeleton, a cell subject to pressure adapts its volume with a certain delay according to the equation

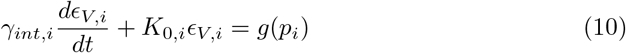

where *γ_int,i_* is a lumped parameter expressing the relaxation behavior after an imposed change of the pressure. It is related to the relaxation time by *γ_int,i_* = *K_i_T_rel_* for a single cell (an analogous argument applies to the whole spheroid). The relaxation period may range from several seconds or minutes up to hours, depending on how long the stress has been applied [12,42,55]. This is related to both intracellular and intercellular reorganizations. In our simulations, we assume 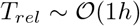 for viable cells motivated by observations of relaxation times in compression experiments [43]. For *K_i_* = *K*_0,*i*_ we have *g*(*p_i_*) = *p_i_*, while in case of a dependency as by Equation 1 it is *g*(*p_i_*) = *K*_0,*i*_ log(*p_i_*/*K*_0,*i*_ + 1).

#### Measures for stress and pressure

The external pressure *p_i_* on a cell *i* is derived from the viral stress and given by:

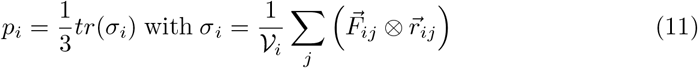

being the stress tensor quantifying the stresses cell *i* experiences subject to contact forces 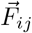 with other cells *j* [20]. Here, 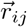 is the vector pointing from the center of cell *i* to the cell *j* with 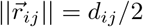 and 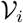 is the *sampling* volume which can be taken as the cell volume. The stress tensor can be diagonalized in order to find the principal direction of stress.

#### Cell growth, mitosis, and lysis

Our basic model assumes constant growth rate during the cell cycle and updates the volume *V_ref,i_* of cell *i* in time as

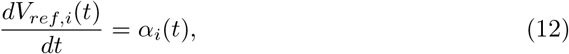

where *α_i_*(*t*) is the growth rate. We studied both, a constant volume growth rate (*α_i_*(*t*) = *C*_1_) and an exponentially increasing cell volume mimicked by *α_i_*(*t*) = *C*_2_ × *V_ref,i_*(*t*) [29–32]. The cell cycle times in both cases are equal for *C*_2_ = log 2 × *C*_1_/*V*_0,*i*_. However, on the time scale (several days) of growth considered here, growth rate variations on time scales of an hour turned out to be negligible. After a cell has doubled its reference volume, it splits into to spherical cells (see the Appendix for more information).

Cells dying either by apoptosis or necrosis eventually undergo lysis. During lysis they gradually shrink. In experiment I the necrotic core appeared very solid like, indicating that the water was drained as a consequence of the high pressure. We mimic the lysing process by setting first *V_ref,i_* → *ϕV_ref,i_* after necrosis, where *ϕ* is the volumetric solid mass fraction. The cell volume change rate is mimicked by Equation 10 and controlled by *γ_int_*. We assumed that lysis times *T_lys_* have a physiological range of 5h to 15 days [26], and we set *γ_int_* ∼ *KT_lys_* in Equation 10 during lysis.

### Deformable Cell Model (DCM)

Agent-based models permitting large deformations and representing cell shape explicitly are generally called Deformable Cell Models (DCMs) [20,56–58,71]. In a basic DCM the cell surface is discretized with nodes which are connected by viscoelastic elements. Nodes and their connecting elements represent a flexible scaffolding structure. The discretization can be extended to the entire cell cytoplasm and even organelles be represented, yet here we regard the cell interior as a homogeneous matter. The nodes at the boundary form a triangulated structure, accounting for the mechanical response of the membrane and cortical cytoskeleton. The total force on each node consists of cell-cell interaction and intracellular interaction forces, the latter describing membrane and cortex mechanical behavior, and cell volumetric compressibility.

The basic equations of motion in DCM is formally the same as for the center-based model (Equation 4), but is now applied to each node *i* of a cell^2^:

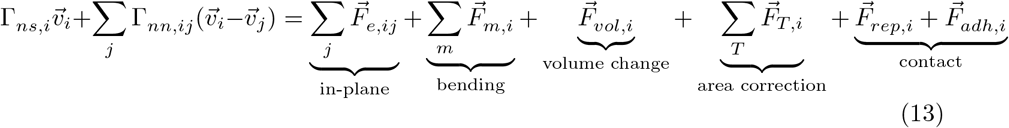

with the matrices Γ_*ns*_ and Γ_*nn*_ representing node-substrate friction and node-node friction, respectively. 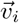 denotes the velocity of node *i*. The first and the 2nd term on the rhs represent the in-plane elastic forces and bending force, the third term on the rhs a volume force controlled by the cell compressibility. The fourth term is a force that avoids excessive triangle distortion. The two last terms 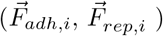 describe the adhesion and repulsion forces on the local surface element in presence of nearby objects as e.g. another cell or the capsule in experiment I (details see Appendix Deformable Cell Model: friction terms and forces). Different from CBMs, the cell bodies in contact do not overlap and therefore triangles belonging to different cells will be repelled upon approaching each other. For consistency with the CBM we chose the model components of the DCM such that cells are inherently isotropic. As the DCM directly represents cell compartments, the range of its parameters can readily be determined (Table 1, for further details, see Appendix).

### Calibration of the CBM contact forces using DCM

During the process of compression, cells rearrange and deform to a closer packing. As discussed above, common models to model the interactions between cells (such as Hertz, JKR, extended Hertz, Lennard-Jones, etc.) base on pair-wise interaction force calculations and do not take into account the effect of volume compression emerging from the simultaneous interaction of many cells [20,48]. In simulations using these interaction force models, the apparent volume (as seen in the simulation) that the spheroid occupies upon strong compression, may become much smaller than consistent with the material parameters; even incompressible cells having Poisson ratio *ν* = 0.5 reduce their volume [20,59]. Simulations of spheroid growth in a capsule performed with an uncalibrated model result in an unrealistic capsule dilatation (see Figure 12).

The deformable cell model (DCM) does not suffer from such shortcomings, but is not amenable to the amount of cells observed in experiments I and II in reasonable computing time on standard desktop computers. For this reason we here chose a hybrid strategy: we corrected the interaction force in the CBM based upon numerical compression experiments performed with the DCM, and used the so calibrated CBM to perform simulations reminiscent of virtual computer experiments in the experimental settings I and II (Figure 8).

**Fig 8.**
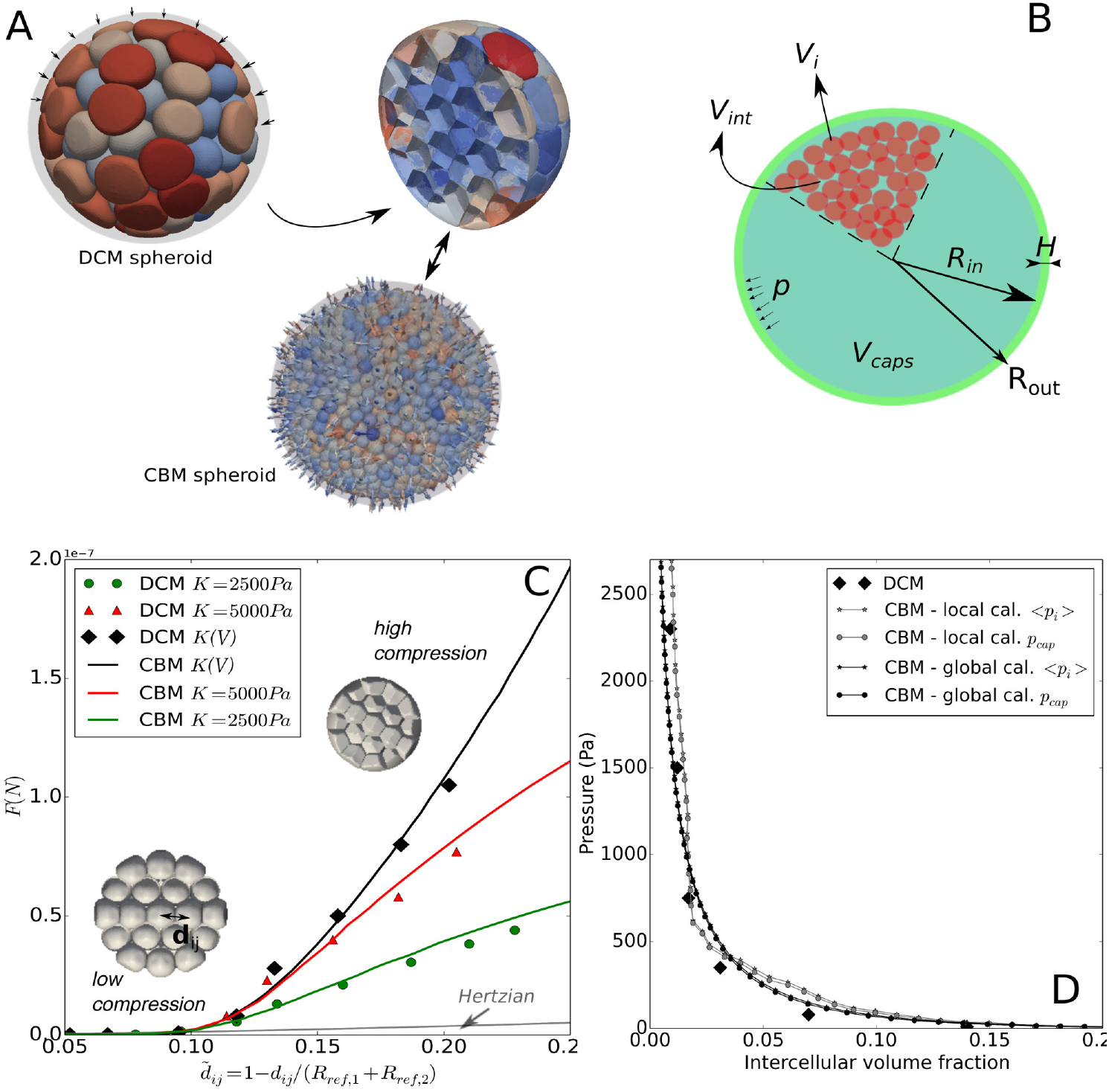
(A) Cartoon illustrating the compression experiment using deformable cells in a capsule to calibrate the center-based model. (A, bottom) Equivalent compression experiment using the center-based model with indication of the maximal principal stress directions of the cells in the capsule during compression using Equation (11). (B) Cartoon showing the volume compartments *V_i_, V_int_* and *V_caps_* in a capsule with thickness *H*. (C) Average contact force vs. 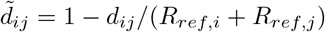 for different *K* values simulated using DCM (diamonds), and CBM with corrected Hertz contact force (full colored lines) replacing *E* by 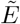, see Equation 14. *d_ij_* is the distance between the centers of cells *i* and *j, R_ref,k_* the radius of a free cell *k* ∈ {*i,j*}. The modified Hertz force shows the same evolution as the force in the DCM, while an uncorrected Hertz force (gray line, Equation 6) strongly underestimates the interaction force for strong volumetric compression. (D) Pressure curves during compression of the spheroid as a function of the inter-cellular volume fraction simulated with the DCM and the CBM with modified Hertz force. The pressure for CBM was computed using both the capsule pressure and average virial stress per cell calculated from Equation (11). A representative movie (Video 3) of these simulations is provided in the Supplementary Material)

In order to estimate the repulsive contact forces in case of many cell contacts, we have constructed a DCM spheroid computer experiment with ∼ 400 cells initially positioned in a closest sphere packing. In this computer experiment, the outer cells were then pushed towards the spheroid center quasi-statically to avoid friction effects, using a shrinking large hollow rigid sphere encompassing the cells (see Figure 8A). All cells have the same size but taking into account a moderate variable cells size were found to not affect the results significantly. Interestingly we observed in the calibration simulations, that cell shape of isotropic cells in the calibration compression simulations with the deformable cell model appear distorted near the capsule border in agreement with the shapes one would infer from the position of the cell nuclei in the capsule experiments [21].

#### “Local” calibration approach, needed for experiment II

For the DCM simulations we adopted *E_cor_* ≈ 2400 Pa, *h_cor_* ≈ 100 nm and *ν_cor_* ≈ 0.5 [37] as fixed elastic properties of the cortex. The cortical stiffness *E_cor_h_cor_* = 0.24 mN/m, is close to values deduced from other experiments performed on fibroblasts [60]. As the cell compression modulus *K* maybe variable and further plays a significant role in this work, we constructed the calibration method such that it works for different values of *K*.

During the simulated DCM compression experiment (Figure 8A) we “measure” all the contact forces between a bulk cell *i* and the surrounding cells *j* in our simulation, which gives us the force, pressure and volumes change on that cell, as a function of their relative positions, 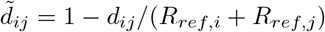. The distance *d_ij_* is computed as the length of the vector connecting the two center of masses of the two cells *i, j. R_ref,k_* is computed as 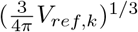, with *k* = *i, j*. The average contact force of the central cell *i* with its neighbors *j* as a function of the cell-to-cell average distance 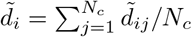 (*N_c_* = number of contacts) is depicted in Figure 8C, for *K* = 2500 Pa, 5000 Pa, and a variable *K* = *K*_0_(*V*) using *K*_0_ = 5000Pa due to strain hardening (see section Cell volume and compressibility). Overall we find that this contact force curve still can be characterized as initial Hertzian contact for 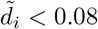, but is after a transition zone followed by a steep increase 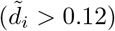. The first part in this curve is largely determined by the mechanical properties of the cortex and the changing contact area of the cells, whereas the behavior at larger compression is determined by the bulk modulus of the cells.

We have developed a CBM calibration approach where we keep the original Hertz contact law (Equation 6) but replaced the Young modulus Ei by an apparent contact stiffness *E_i_* (i.e., *E_i_* → *E_i_*) of the cells as they get nearer to each other. In other words, 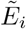 gradually increases in Equation 6 as the cells get more packed, based on the reasoning that indenting a piece of material with another object gets more difficult when confined. The total strain of the cell is composed of a deformation of the cortex largely determined by the apparent stiffness 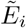 and the volumetric compression determined by *K_i_*. The volume (and radii) of the cells are adapted using Equation 10. It is important to stress here that 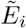 only reflects the contact stiffness of the cell through Equation 6, while the bulk modulus (Equation 8) is determined by the original cell Young’s modulus *E_i_*.

To take into account the limited cell volume compressibility in a pairwise cell-cell interaction force, we fitted 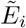 by a function that depends on the local average distance 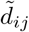 for a bulk (i.e., interior) cell in the simulated experiment in Figure 8A:

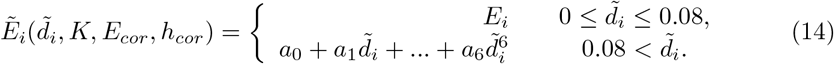

Here, the *a_k_* with *k* ∈ [0, 6] are fit constants (see Appendix). They are calibrated such that the function is monotonically increasing and results in an optimal fit to the average force a cell *i* experiences upon compression of the cell aggregate (see Figure 8A) as function of the distance between the center of cell *i* and its neighboring cells *j* in the DCM simulations (see Figure 8C). The higher the compression, the higher gets the contact stiffness, so that at strong compression, the contact forces only result in a very small increase of indention, yet the cell volume decreases (Equation 10).

At the point of confluence when outer cells touch the capsule wall, the DCM cells exert a total interaction force *F_cap_* = Σ_*i*_ *F_cap,i_* on the capsule wall. The capsule pressure was then computed by *p_cap_* = *F_cap_/A_cap_* where *A_cap_* is the inner surface area of the capsule. On the other hand we defined the intercellular volume fraction, as *ϵ_int_* = *V_int_/V_cap_* (see Figure 8C). Here *V_int_* = *V_cap_* – Σ_*i*_ *V_cell,i_* is the volume of the space in between the cells, *V_cap_* is the total capsule volume. We then compared for the DCM simulations and calibrated CBM the resulting pressure versus intercellular volume fractions. These curves do not match exactly, but follow each other closely (Figure 8D).

We further complemented this study by pursuing a “global” approach where we estimated the forces and pressure exerted by the MCS on the capsule as a function of the total intercellular space fraction occupied by cells within the elastic capsule (see Appendix, section “Global” calibration approach, only valid for confined spheroid), obtaining the same results. Both calibration approaches can be used for arbitrary values of *K*.

### Elastic Capsule Model

The capsule is made of an quasi-incompressible alginate gel exhibiting a strain hardening behavior. The stress-strains relationship was measured in a stretching experiment of an thin alginate cylinder. Strain hardening behavior was observed for strains > 15%. In case of a thick walled capsule, the expansion strain is low and hence linear elasticity can be applied. We refer to the hollow sphere example as described in [61] to compute the radial displacement of the capsule from the internal pressure. If on the other hand the capsule has a thin wall, strains can become large, and the linear elasticity hypothesis fails. For this case, in line with ref. [21] the original young modulus is modified instead of employing nonlinear elasticity theory. The nonlinear relationship in stress and strain (*ϵ_cap_*) was phenomenologically characterized in ref. [21]:

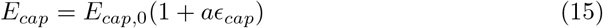

where *ϵ_cap_* is the strain and *a* = 1.5 to obtain an optimal fit with the experiment.

The capsules have an initial inner and outer radius *R*_*in*,0_ and *R*_*out*,0_ respectively, whereby typically *H* = *R*_*out*,0_ – *R*_*in*,0_ > 0.2*R*_*in*,0_ for thick capsules, *H* being the capsule thickness. The pressure difference along the capsule wall can be related to the change in radii by [21]:

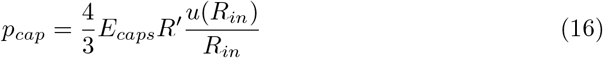

Where *E_caps_* is the Young modulus of the capsule material, *R_out_* is the outer radius, and *u*(*R_in_*) = *R_in_* – *R*_*in*,0_ is the displacement at the outer radius. Furthermore, 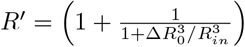, in which the outer radius is related to the inner radius *R_in_* by 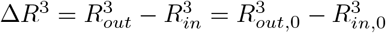, assuming incompressibility of the elastic shell. To simulate the radius evolution of the capsule, one computes pressure *p_cap_* by dividing the sum of all contact forces of the cells with the capsule by the actual inner surface area. Taking into account the damping by the alginate material, we arrive at an ODE, formally similar to Equation 10:

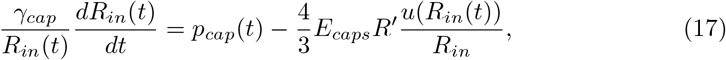

with a lumped material damping parameter *γ_cap_*. It was shown in [43] that the viscosity of the capsule material is low and does not influence the much slower dynamics of the spheroid. Accordingly, in our model *γ_cap_* was chosen low to reflects the material’s ability to rapidly adapt to a change in spheroid radius while not affecting the slow growth dynamics.

### Model setup and parameter determination

We here explain the determination of the mechanical model parameters starting from the thin capsule experiment. A large fraction of the parameters are fixed from direct observations or published references, see Table 2 for more details.

Within parameter sensitivity analysis simulations the parameters that could not be fixed by experimental observations, were varied within their physiological ranges to study their impact on the simulation results. Some parameters turned out to only negligibly affect the simulations results, see section Model parameter and algorithm sensitivity.

As the simulation time was too long to determine the parameters within their physiological ranges based on a maximization of a likelihood function, or to perform a parameter identifiability analysis, we identified plausible parameters by a two-step procedure.

We first determined those model parameters that determine the simulated growth behavior in case of free growth by comparison to the experimental data for CT26 in experiment I. In the next step the parameters relevant for the specific experiment were fixed. After this, two remaining parameters, namely *K* and *T_lys_* were calibrated by the thin capsule simulations, yielding a model without a growth rate adaptation (see section Cell-specific parameters *K* and *T_lys_* during stress conditions).

Each simulation result was compared to the experimentally observed spheroid diameter of the growing spheroid prior to confluence, and the slope of the residual growth curves after 48h, thereby retaining the parameters that are physically plausible and can best explain the data at the same time.

#### Cell-specific parameters {*P*_*C=j*_} to obtain the initial spheroid configuration and free growth

Starting from the calibrated model (step 1), a single run was performed with a small aggregate of 10 CBM cells, all at the beginning of their cell cycle, to grow a spheroid up to the size of *R* = 100 *μ*m, which corresponds to the size before confluence, see Figure 2B. A cell cycle time of *T_cyc_* = 17 was assigned to each of the cells as this matches the experimental observation. Cells increased their radius from ∼ 6 *μ*m until their radius reached the division size (7.5*μ*m). After each cell division, a new cell cycle time was assigned to each of the daughter cells, randomly chosen from a Gaussian distribution with 〈*T_cyc_*〉 = 17*h* and standard deviation of ±10/%. The intrinsic free growth cell cycle time defines the growth rate *α*_0_ = 1/*T_cyc_*.

The cell-cell adhesion energy *W* determines how close the cells approach each other in aggregates not subject to compression by external forces, and has been chosen such that the area density, measured in a cryosection of width 10 *μ*m of the resulting spheroid with *R* = 100*μ*m, matches that of the experiments (∼ 0.85/100*μ*m^2^) [21]. In these simulations the cells have a fixed Young’s modulus of *E* ∼ 450 Pa and a cell motility coefficient *D* of 10^−16^ m^2^/s [18]. The compression modulus was here set to *K* = 5 kPa inferred as an average from values reported in literature, see Table 2. For MCS grown in absence of external stress, *K*, if varied in the range of experimentally observed values, had no significant effect on the growth simulation results.

The physical parameters responsible for the inter-and intracellular friction are in the CBM represented by *γ_int_, γ_cc_*,⊥, *γ_cc_*,∥ ∈ {*P_C=j_*}. Mechanical relaxation time of spheroids compressed over a longer time period indicate relaxation times of 1 to 5 hours in experiments [42,43]. We have calibrated the friction parameters in the model from a relaxation experiment starting from a compressed spheroid (see Figure 8A) such that *T_rel_* ∼ 2h, lying well in the reported range [1*h*, 5*h*], was obtained using as observable the spheroid size as function of time. The calibrated coefficients correspond to those found in [40,41].

The parameter set as determined above resulted in a good agreement for free growth simulation with data from experiment I. The model robustness was finally tested by varying these parameters to see how they affected the simulation results of the thin capsule (see Appendix Model parameter and algorithm sensitivity).

#### Experiment specific parameters {*P_EXP_*}

Here, we determined the parameters that are exclusively related to the experiments. See Table 2 for an overview.

**Experiment I**: From the data for the capsule radius at which the curve is in the transition stage *T*_1_ to *T*_2_ (Figure 9, *t* = 1*d*) and using Equation 16, a pressure of *p_th_* ∼ 1500 Pa could be inferred (Figure 4C), at which bulk (interior) cells further away from the border than λ_*I*_ are experimentally observed to become necrotic. To express the variability in the cells’ response on pressure we chose *p_th_* from a Gaussian distribution with mean 1500 Pa and standard deviation of 150Pa (10%) in all simulations. A variation of ±300 Pa on the mean value reduced the agreement with data in all simulations. The rim thickness λ_*I*_ within which the cells remain viable is fixed during the simulations as it did not change during the experiment. Notice however, that the value of λ_*I*_ does not explain the MCS expansion speed that differs for the thick capsule from that for the thin capsule, as it is demonstrated below (Figure 9A). We further assumed that cell-capsule friction coefficients *γ_c,cap_* are similar to those of cell-cell friction. However, the simulation results are robust with respect to wide variations on friction parameters, see Appendix Model parameter and algorithm sensitivity. The elastic properties of the capsule are fixed to the values measured in [21].

**Fig 9.**
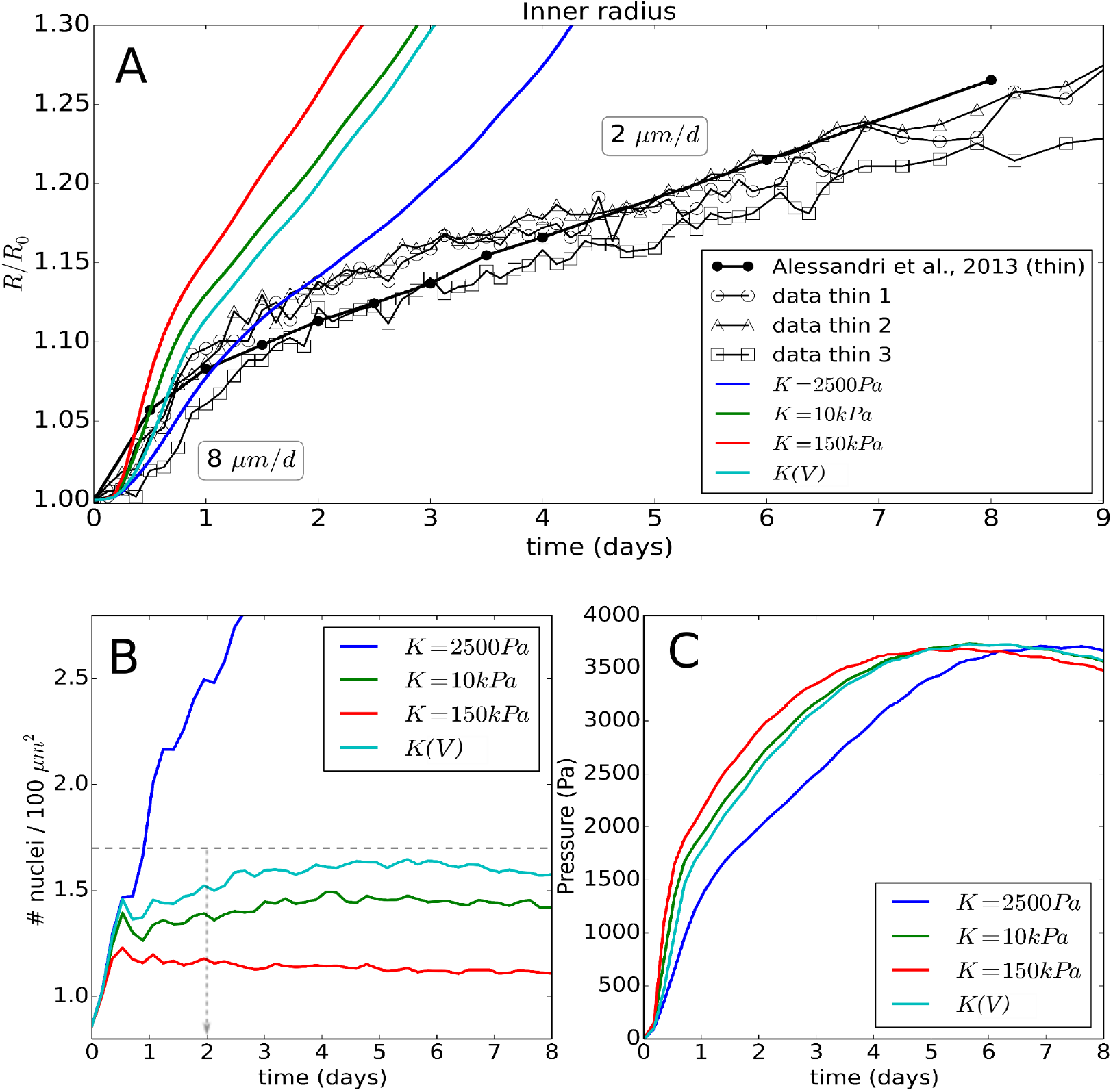
(A) Time evolution of the radius of the thin capsule, shown for the experimental data and the simulation using Model I, with parameter variation on the individual cell compressibility (*K*(*V*) means strain hardening). (B) Time evolution of the simulated cell density. The dashed horizontal line indicates the experimentally observed cell density at 48h. (C) Pressure in the capsule versus time.

**Experiment II:** In Experiment II, λ_*II*_ was calibrated to match the growth rate kinetics of the spheroid in the absence of dextran (see EII.OII). Contrary to Experiment I, after adding external mechanical stress via dextran, no increase of necrosis was observed (EII.0II). This was formally captured by setting *p_th_* → ∞ to in the model. The magnitude of the osmotic forces to obtain the desired bulk spheroid pressure was computed from Equation 3, fixed for each experiment.

#### Cell-specific parameters *K* and *T_lys_* during stress conditions

In the next step the compression modulus and the cell specific lysis time have been specified. To acquire the most realistic parameters within their physiological range, we consider the spheroid growth in the capsule, first with the constant growth rate *α*_0_ of the cells as determined from free spheroid growth in Experiment I.

#### Compression modulus of the cells

The compression modulus of the cells influences the volumetric strain and hence through Equation 10 the growth rate *α*. First we tested the hypothesis that *K* remains constant during the experiment, varying *K* in the range *K* ∈ [2.5 kPa, 150 kPa] in simulations for Experiment I. *K* ∼ 2.5 kPa has been measured for quasi uncompressed L929 fibroblasts [37], *K* ∼ 10kPa for compressed CT26 cells [12].

Simulations with *K* = 10 kPa resulted in a cell density increase at 48*h* by only a factor of 1.5, while experimentally a factor of two is observed (Figure 9B), suggesting that this value of *K* is too high. Moreover, a significant overestimation of both the initial and the residual radial growth could be observed (Figure 9A). We further tested two extremes for *K*. For *K* = 150 kPa the cell density at 48h is now only 1.3 times the original one (Figure 9B), with a largely overestimated initial radial growth. By contrast, for a much smaller value *K* ∼ 2.5 kPa, the cell density is strongly overestimated (increase by 3-fold at 48*h*), hence we reject such low values.

In a next step we tested the consequence of strain hardening (section Cell volume and compressibility, [51–53]). *K*(*V*) can be initially relatively small, leading to a higher overall cell nuclei density (Figure 9B), yet gradually increasing during compression. For an applied pressure of 5 kPa, we find *K*(*V*) = 10 kPa while for an applied pressure of 10kPa we have *K*(*V*) = 15kPa, comparable to the values reported in [12,38]. The simulations with strain stiffening show a better estimation of the cell density at 48*h*.

However, the stiffening alone did not solve the discrepancy between data and model simulation results. It allows a rapid nuclei density increase in a spheroid for low pressure but at the same time leads to higher mechanical resistance with increasing pressure. The capsule pressure generally shows a highly nonlinear behavior with a maximum (Figure 9C). This is typical because the mechanical stiffness of a capsule drops at high dilatation [62] as confirmed in the experiment by the observation of cells sometimes breaking through the capsule at later stages [21].

Note further that all the simulations of the capsule radius upon deformation by the growing MCS with time exhibit a short initial lag, in where the capsule dilatation is small (Figure 9A). In this stage, the spheroid touches the capsule border but cells are mainly pushed inwards, filling up intercellular spaces. This is less visible in the experiment, yet there the exact point of confluence is difficult to determine. After this period, cells are becoming more and more compressed and the mechanical resistance of the spheroid increases.

Overall, these results demonstrate that the proliferating rim with λ_*I*_ = 20*μm*, constant growth rate and neither constant nor strain-dependent growth rate cannot explain the velocity of the growing spheroid in the linear phase, as it is not possible to simultaneously fit the nuclei density and the long-time radius expansion. For any value that would be capable of fitting the nuclei density, the slope of the radius expansion would be too high.

#### Lysis time

In a next step we studied whether incorporating the effect of intrinsic volume loss of necrotic cells due to lysis would lower the radius expansion and establish agreement between model and data. Lysis as defined in ref. [26] induces an irreversible water loss and decrease of cell volume (see section Cell volume and compressibility) limited to the solid volume of the cell. Contrary to *in vivo* experiments, there are no macrophages present to phagocytose the remaining cell bodies, and phagocytosis by neighbor cells is very slow [24]. In line with [26], we studied lysis times *T_lys_* ∈ [5*h*, 14*d*] using Model I. We notice that the shorter *T_lys_*, the more the curves bend off in the beginning. However, because lysis results in more compression and thus gradually leads to stiffer cells, the numerical growth curves largely fail to reproduce the observed linear behavior (see Figure 10). The effect becomes striking at very low lysis times (*T_lys_* = 5*h*). Here, the initial behavior of the spheroid is determined by cells quickly loosing their volume (hence a low resistance against pressure). Further in time, a large stiff core develops which will eventually overcome the mechanical resistance of the thin capsule. Nevertheless, adopting *T_lys_* ≈ 5*d* yields a good agreement with the cell nuclei density at 48h (Figure 10B), which is in line with values found in an in silico model for ductal carcinoma in situ [26] and is relatively close to the apoptosis time found by fitting phenomenological growth laws for spheroids (∼ few days) [12,63]. Note that the lysing cells in the bulk tend to move very slowly towards the center of the spheroid (see the Supplementary Material, Video 2).

**Fig 10.**
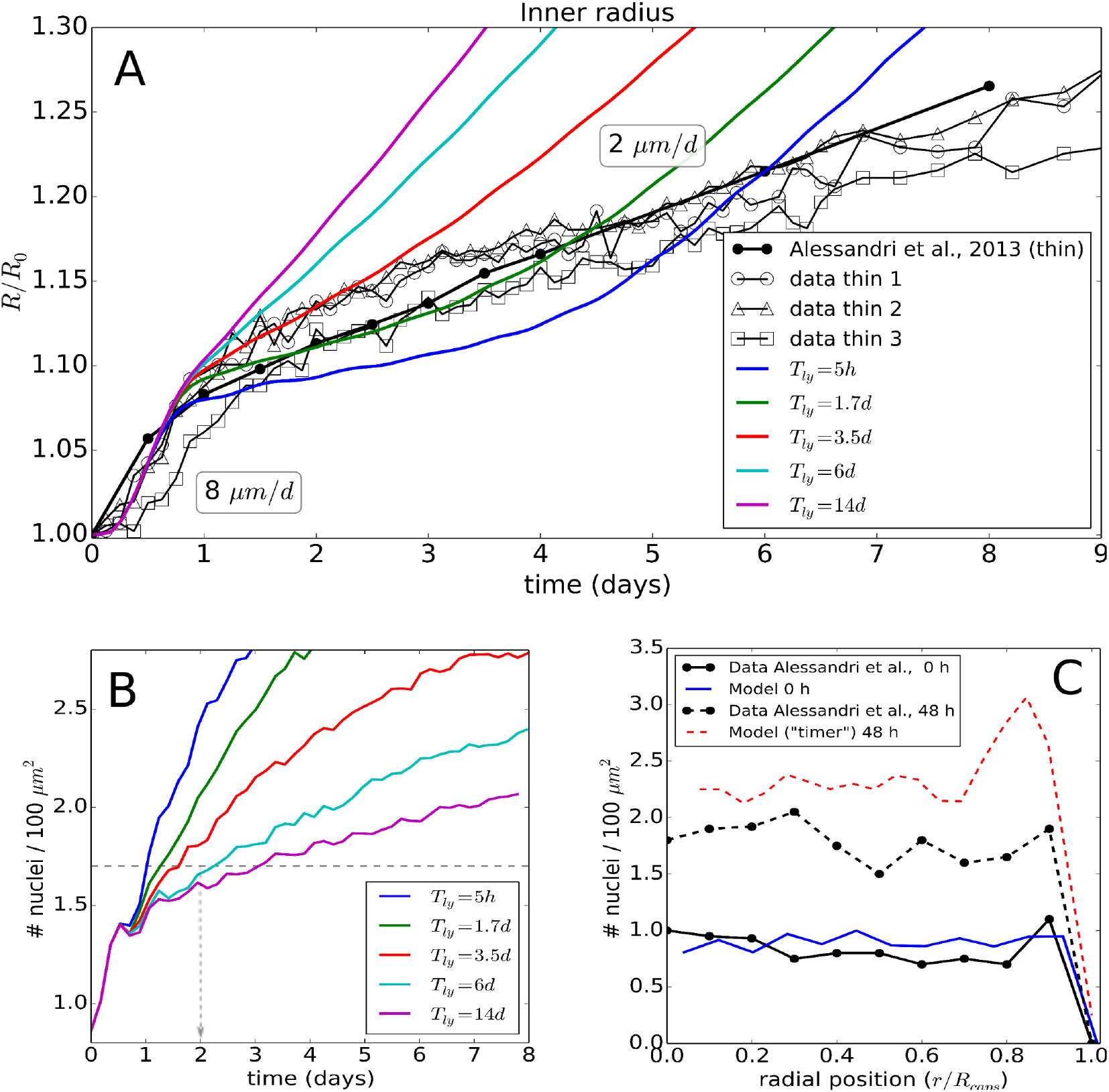
(top) (A) Time evolution of the radius of the thin capsule, shown for the experimental data and simulations using Model I, showing the effect of a parameter variation for the lysis time *T_lys_*. (B) Time evolution of the simulated cell density. (C) Cell density at 48*h* obtained from final model run with optimal parameters, but in which cells divide after a fixed cycle time (“timer”) instead of a fixed size.

#### Non-constant growth rate

Even including lysis it was still not possible to simultaneously fit growth and density curves as improvement of growth kinetics was accompanied by increasing mismatch of density and vice-versa. This prompted us to study non-constant growth rates, decreasing with increasing volumetric strain as explained in the main text.

## Acknowledgements

This work was supported by INSERM PhysCancer (France), ITMO INVADE (France), the EU 7th Framework Programme (Notox), Bundesministerium für Bildung und Forschung (Virtual Liver Network, LiSym, LEBERSIMULATOR), Agence Nationale de la Recherche (iLite). P.N. acknowledges financial support from INCA (grant 2012-1-PL BIO-09-IO) and from the ANR Blanc SVSE5 “Invaders”. We would like to thank M. Delarue for his input. P.N. would like to thank J. Prost for the useful discussions.

## Supplementary Material

### Appendix

Below further technical details on the models and parameters calibration are explained. A flow chart of the model algorithms executed in the simulations is depicted in Figure 14.

#### Center-based model: friction terms and forces

We look at the Equation of motion (4) for cells in more detail. The general form for the friction tensors in Equation (4) reads

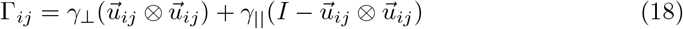

with 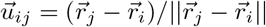, where 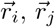 denote the position of the centers of cell *i* and object *j. I* is the 3 × 3 identity matrix, 0 denotes the dyadic product [20]. The individual cell friction coefficients are *γ*⊥ and *γ*∥, respectively perpendicular and parallel to the movement direction. As experiments do not indicate ECM inhomogeneity or anisotropy, the cell-ECM friction matrix is considered to be diagonal. The ECM was not represented explicitly as experimental observations indicated that the ECM fraction is small and approximately homogeneously and isotropically distributed in the intercellular spaces.

As represented in Equation 5 the interaction force resulting from compression, deformation and adhesion can be expressed as a sum of a repulsion force, here represented by a modified Hertz contact force, and an adhesion force. The interaction force acts along the line connecting the centers of two interacting spherical cells.

The micro-motility force of cell *i* is mimicked by a Brownian motion term with zero mean value and are uncorrelated in time:

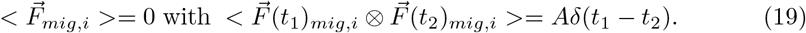

Based on a formal analogy to colloidal particle systems [64], the autocorrelation amplitude of the force as approximated *α* = 2*Dγ*^2^*I, D* being the cell diffusion constant, *γ* the friction coefficient of a cell in the medium and *I* the unity matrix. The scalar *γ* emerges in an isotropic environment, for which *γ*∥ = *γ*⊥(:= *γ*). In an extracellular matrix environment, *α* is largely controlled by the cell itself [18]. The equations of motion (4) do not conserve total momentum due to the micro-motility term as part of the momentum is transferred to the ECM, that is not explicitly modeled here.

The system of Equation 4 was integrated numerically until the simulation time surpasses the total duration of the experiment which is ∼ 10*d*. Equation 4 results in a linear problem with a sparse symmetric matrix, which can be solved efficiently by a Conjugate Gradient method [20,65] and an explicit Euler integration scheme.

#### Deformable Cell Model: friction terms and forces

We recapitulate the basic equation of motion for a DCM denoted in (Equation 13). The individual elements generating the in-plane elastic forces between the surface nodes represented by the 1st terms on the lhs and rhs of Equation 13 are modeled by classical linear spring-damper systems. The force between the nodes captures the elastic response of the shell-like structure including the cortical cytoskeleton of the cell. When the elastic and dissipative components are summed up, one acquires a Kelvin-Voigt element. The vector force between two nodes *i* and *j* reads (see Figure 11B):

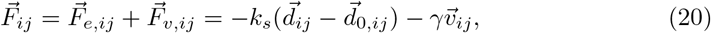

where 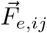 and 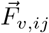 are the elastic and dissipative forces, *k_s_* is the spring constant, *γ* represents the dissipation, 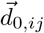, 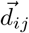 are the initial and actual distance vectors, and 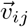 is the relative velocity between the nodes, respectively.

**Fig 11.**
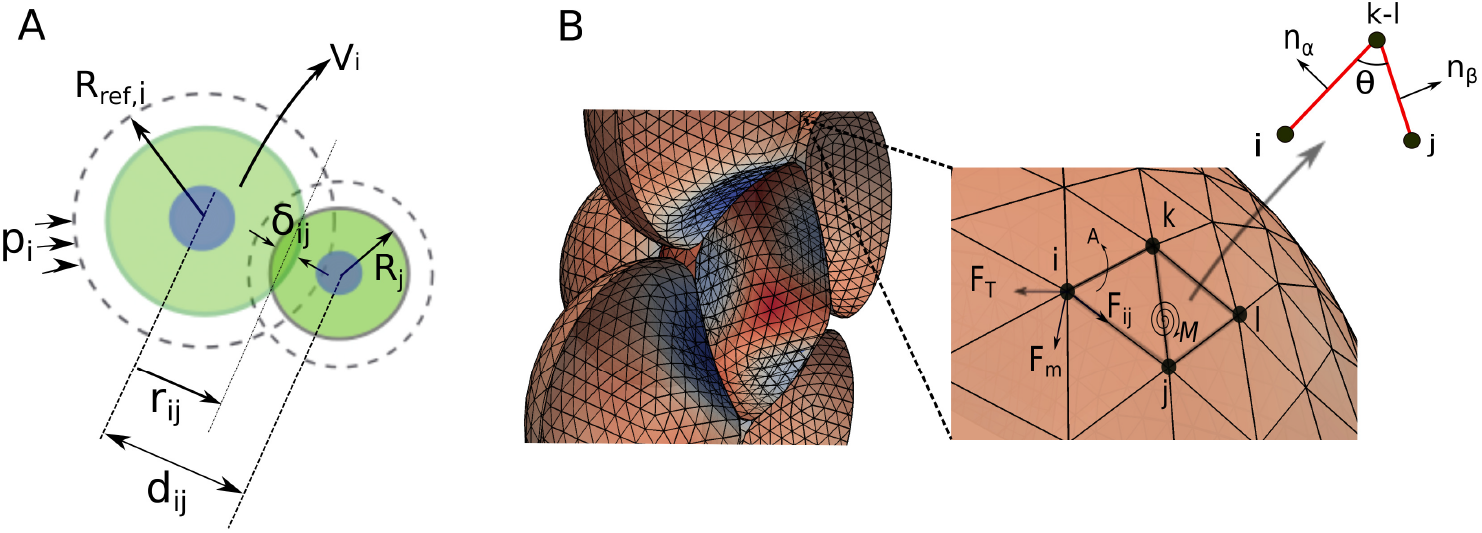
(A) Contact between two cells defining (a) the distance *d_ij_*, virtual overlap *δ_ij_*, reference radius *R_ref,i_*, actual radius *R_i_* and volume *V_i_*. (B) (left) DCM representation of several adjacent cells. (right) Detail of nodal structure building cell surface and depicting the forces that work on them.

The second term in Equation 13 representing the surface bending resistance is incorporated by the rotational resistance of the hinges determined by two adjacent triangles *α* = {*ijk*} and *β* = {*ijl*} (Figure 11B). This defines the bending moment *M*:

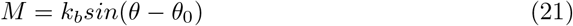

where *k_b_* is the bending constant, and *θ* is determined by the normals to the triangles 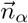,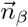, with *θ*_0_ being the angle of spontaneous curvature. *M* can be transformed to an equivalent force system 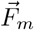 acting on each of the nodes of the triangles [65]. Restoring volume compression / expansion forces 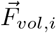 controlled by the bulk compressibility of the cytoplasm are computed from the internal pressure using the volume change and compressibility modulus of the cytoplasm *K*. In our simulations *K* reflects the overall cell compressibility including the cortex, as in its physiological range of elasticity, the cortex contributes little to the overall bulk modulus of the cell, i.e. *K* ≫ *E_cor_ h_cor_/R_cell_* ([37], own test runs). In analogy to Equation 8 the pressure applied to the cell is therefore approximately given by:

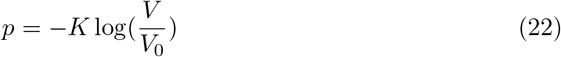

The forces 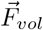 on the nodes are perpendicular to the cell surface and can be obtained by multiplying the pressure with the surface area assigned to each node (each node has 6 adjoining triangles which each count for 1/3 of the surface area).

During the simulations, large variations in area of the triangles in the network can cause numerical artifacts. These can be avoided by adding a force *F_T_*, which is proportional to the area expansion of the individual triangles [66,67]:

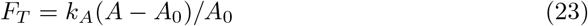

Here, *α*_0_ and *α* are the initial and the current areas of the triangle, and *k_A_* is the area compression stiffness. The forces 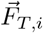 are summed over all triangles of the cell and transferred to the nodes in the direction perpendicular to the opposite vertex edge of that node leading to increase (or decrease) the area of one triangle at the expense of a decrease (or increase) of its neighbor triangles. We chose *k_A_* such that the influence on the cortex elasticity is minimal, yet triangle deformations are minimized. Note, these forces do not induce shear elastic effects in the network.

Finally, the interactions between neighboring cells are accounted for by introducing repulsive forces 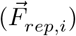 and the adhesive forces 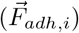 between nodes belonging to different cells. The interaction forces were obtained as detailed in [67]. The model allows simulating the interaction between two arbitrarily shaped triangulated bodies but also between a triangulated body and non-triangulated simple geometric objects as spheres and flat substrates.

Importantly, the parameters of the spring network in Equation 20 can be related to macroscopic elastic constants, approximating the cell cortex by a thin shell. For the six fold symmetric triangulated lattice on the cell surface, the linear spring constant *k_s_* can be computed from the Young modulus *E_cor_* of the cortex with thickness *h_cor_* by [54,68,69]:

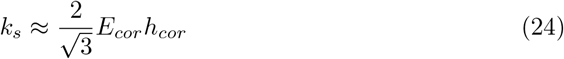

Similarly, the bending stiffness of the cortex can be approximated by

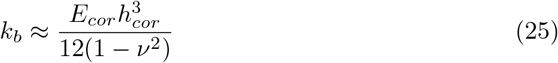

where *ν* is the Poisson ratio (= 0.3 for an equilateral 2D network of linear springs).

#### “Global” calibration approach, only valid for confined spheroid

The relationship between forces, inter-cellular volume fraction and pressure established in section “Local” calibration approach, needed for experiment II can also be computed following a simpler, global approach in case of a confined volume as in experiment I. If compression rates are sufficiently slow as is the case in that experiment, the cells in the spheroid can reorganize, distributing the stress isotropically and homogeneously over the cells in the spheroid. For the intercellular spaces in the capsule the *global* intercellular volume fraction *ϵ_int_* and the cellular fraction *ϵ_cells_* are related by *ϵ_int_* + *ϵ_cells_* = 1 with *ϵ_cells_* = *V_cells_/V_caps_*. We can now use *ϵ_cells_* to re-parameterize Equation 14 assuming cells are homogeneously distributed over the spheroid during compression (compare Figure 8A). The apparent contact stiffness 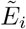 in that case increases for all cells equally with *ϵ_cells_* being the equivalent of 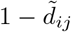 in the previous approach.

With both different calibration methods, local and global, the CBM simulation results closely follow the DCM curve (Figure 8D; Figure 12). However, the curve using the global calibration method is smoother than for the local calibration method as it represents an average over all cells thus disregarding local fluctuations. For high pressures (*p* > 2kPa), both curves become nearly parallel. On the other hand, as the local approach does not require the existence of an enclosed (capsule) volume it can be used more generally as a cell-cell interaction force upon volumetric compression in many configurations as in experiment II.

**Fig 12.**
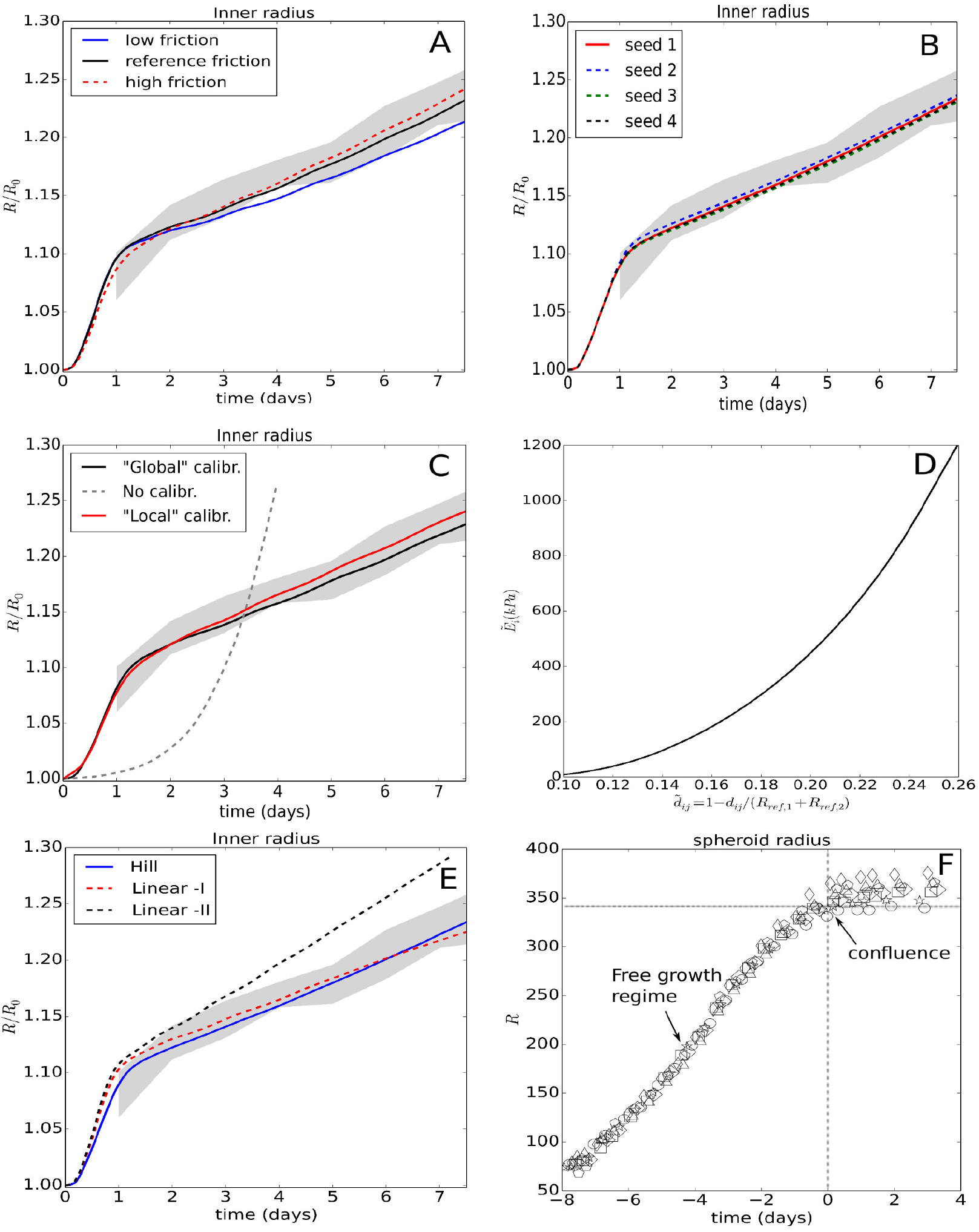
(A) Dependence of simulation results of the thin capsule on a variation on the friction coefficients. (B) Comparison of 4 simulations with different random seed. (C) Comparison of the global and local calibration approach for a growing spheroid in the capsule. (D) Plot of the function 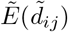 used for cell contact stiffening upon spheroid compression. (E) Simulated thin capsule dilatation using for cell cycle progression: the proposed Hill function, the linear function (“Linear-I”) as depicted in Figure 2 (which fails to explain the data in experiment II), and a linear function (“Linear-II”) optimized to match the CT26 spheroid expansion in Experiment II. (F) Data plots of MCS growth in a thick large capsule (∼ 400 *μ*m) growth. The stress-dominated growth regime is too short to identify the stress-response.

We finally point out that no explicit representation of ECM has been considered in the model based on the observations the ECM fraction is rather small (personal communication) and approximately homogeneously and isotropically distributed in the intercellular spaces.

#### Model parameter and algorithm sensitivity

We also studied the potential influence of parameters calibrated for the free growth conditions which could not be directly inferred from the experiments. For the friction terms, the cell-cell friction, cell-ECM friction and cell-capsule friction were varied between 1 % (“low”) and 200 % (“high”) of their reference values. The effect is shown in Figure 12A and indicates that even for low friction coefficients the results remain largely unaffected. For the cell-cell adhesion energy *W* and cell motility coefficients *D*, which were varied between 10 % and 500 % [23], we did not observe any significant changes (results not shown). The strength of cell-cell adhesion has been shown to play a role in detachment, but to be of minor importance for multicellular systems under compression (e.g. [48]). Note, that also in MCSs growing in absence of externally applied forces cells are moderately compressed [46].

In conceptual analogy to experimental statistical procedures, we have performed growth expansion simulations in the thin capsule with the optimal parameter set but four different random seeds for the cell-specific cycle time, Young modulus and necrotic pressure threshold to test the effect of stochasticity (see Table 2, Figure 12B). Even after more than *γ* days of simulation, only very little mutual differences can be observed, while the slopes of the curves are the same. This can be attributed to self-averaging effects such that the variations on the level of individual cells cancel out at the population level (e.g. ref. [24]).

#### Comparison of calibration methods

For the local calibration procedure, we used the following constants in Equation 14 assuming strain hardening: *α*_0_ = 0.4454 · 10^5^, *α*_1_ = 2.347 · 10^5^, *α*_2_ = −7.918 · 10^6^, *α*_3_ = 6.615 · 10^8^, *α*_4_ = −1.206 · 10^9^, *α*_5_ = −3.091 · 10^9^, *α*_6_ = 1.1239 · 10^10^, yielding a force cell - intercellular distance curve that matches well with the one obtained from the deformable cell simulations experiment (Figure 8). The function for 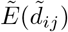 is monotonically increasing (Figure 12D). For the global calibration approach, we obtained a similar curve. In test simulations for the MCS growth in the thin capsule using contact inhibited growth for both local and global calibration, we found only a mutual variation of 5 – 7%, see Figure 12C. On the other hand, if no calibration was assumed, a largely unrealistic capsule dilation was obtained (Figure 12C), and a consistent relation between pressure and cell density could not be established.

#### Influence of cell division algorithms

As far as either the volume or the duration of the cell in the cell cycle have passed a threshold value, we replace the mother cell by two daughter cells which are placed very close to each other [7,40]. This algorithm is different from the approach where cells slowly deform into dumbbells in mitosis phase before splitting into two daughter cells [48,70]. In our model two daughter cells are created next to each other instantaneously. During the mitosis period (which takes about 1.5h) the daughter cells do not grow. However, the two newly created cells can generate short-time artificial pressure peaks which disappear during the division time course due to small local re-arrangements. To test the impact of the pressure peak formation on our growth simulation results we implemented an smoothing algorithm that reduces these peaks by ensuring that (1) the mother cell divides in the direction of the least stress as derived from the local stress tensor (see Methods, section Measures for stress and pressure), and (2) a local energy minimum is sought by varying the distance between the daughter cells and computing the interactions with the other cells. While the smoothing algorithm reduces the short-time pressure peaks, we did not see significant differences in the results compared to simulations where this algorithm had been dropped.

#### Cell deformation and pressure distribution during in a compressed spheroid in DCM

The DCM simulations of a small spheroid compression experiment show that the cells have a flattened shape at the border of the capsule, see Figure 13. In the CBM this is only implicitly captured, by looking at the principal stresses (indicated in Figure 8A by arrows) that can be computed from the stress tensor Equation 11. One observes that the direction of maximal stress points radially to the border cells, while minimal stress direction points tangential to the capsule wall.

**Fig 13.**
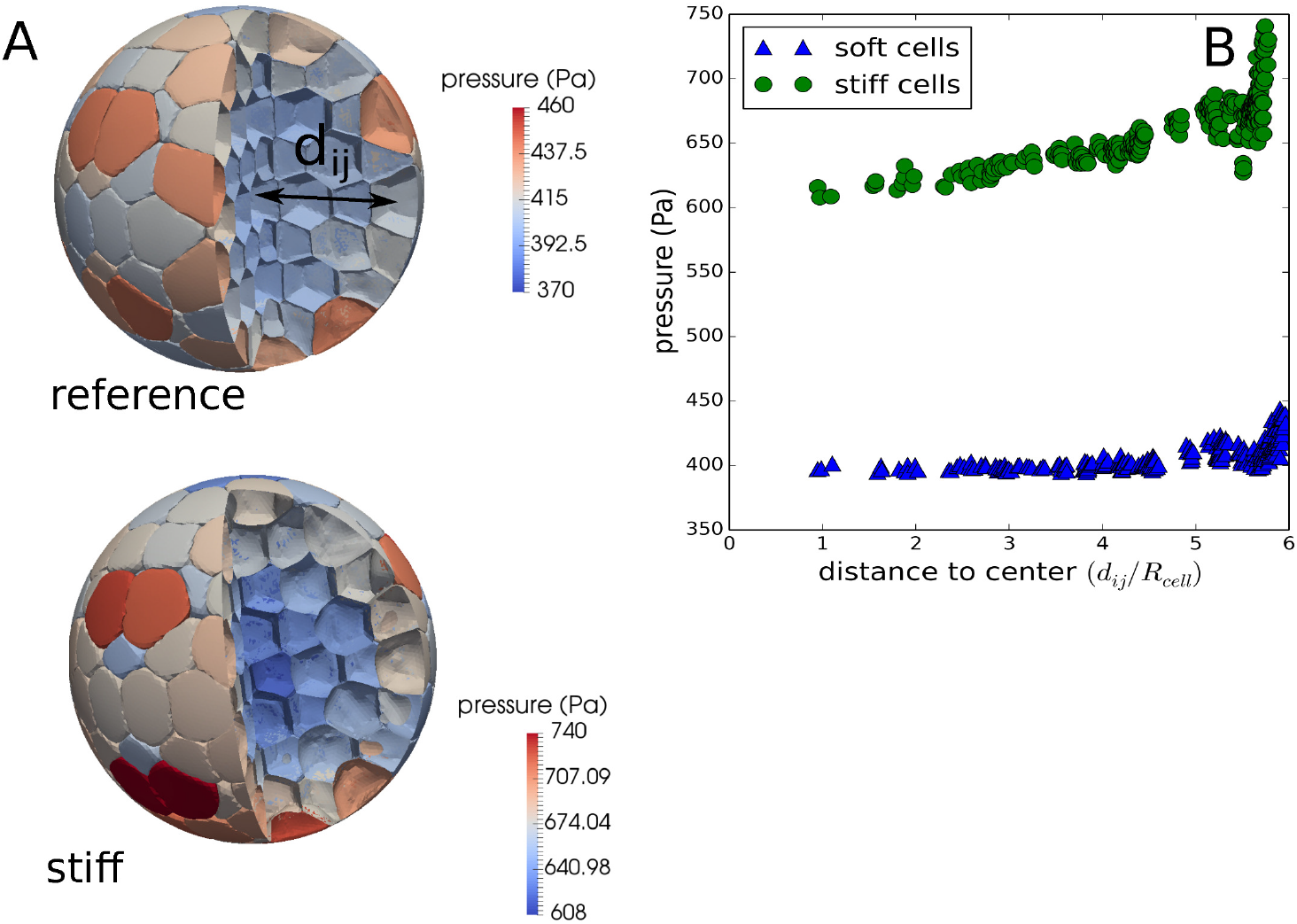
(A) Simulation snapshots of DCM cells within a scaled capsule model, for the cases of cells with a reference cortex stiffness (top) and a “stiff” cortex stiffness (bottom). The coloring is according to pressure (B) Internal cell pressure for deformable cells in a shrunk capsule for nominal cells and stiff cells, as function of distance to the capsule center. The stiff cells show a higher variability in pressure if moving away from the center. Notice that like in the calibration simulations we use cells of equal volume prior to compression but the method can equally be applied to any prior volume distribution.

**Fig 14.**
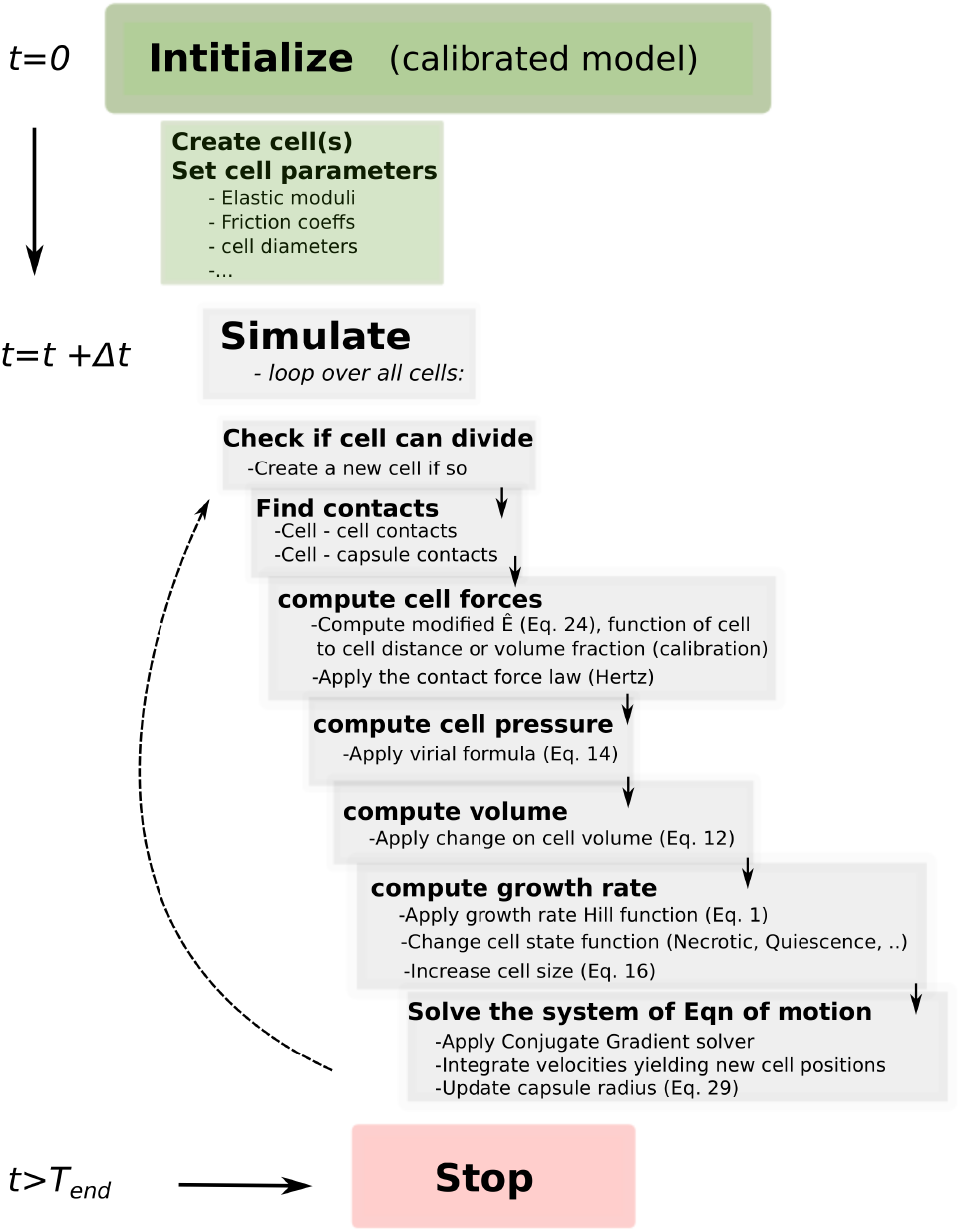
Model flow chart.

We further considered whether the apparent boundary effect (EI.OIII) could be attributed purely to mechanical effects. For this, we used a spheroid compression experiment with a scaled capsule system using 400 (quiescent) DCM cells with different cortex properties (i.e. cells that have the reference Ecor and cells with 10 times this value). It is shown in Figure 13 that there can be a small mechanical effect in the case for a “high” stiffness of the cortex, as the simulations show that the cells near the boundary acquire higher pressures as compared to the bulk cells and a weak gradient from the center to the spheroid edge can be observed. This can be attributed to arching effects, the phenomenon where outer layers of cells bear more stress compared to inner layers. The effect increases with increasing cortex stiffness. On the other hand, reference parametrized cells spread out more easily, diminishing the pressure differences. To investigate the boundary mechanics in a more realistic system with dividing cells, the DCM could be extended with the capability to mimic mitosis. In our *simple* compression experiment with cells having estimated cortex properties, the boundary effect appears acceptable.

## Videos

**S1 Video. CT26_free_growth.avi** shows the simulated evolution of pressure a free growing CT26 spheroid. Note that a gradient in cell pressure gradually builds up from the center to the border of the spheroid.

**S2 Video. CT26_spheroid_capsule.avi** shows the simulated evolution of pressure and cell volume of the CT26 spheroid growing in a thin capsule. The pressure increases gradually but remains approximately uniform over the spheroid.

**S3 Video. DCM_spheroid_compression.avi** shows the simulation of a compression experiment of a spheroid in a capsule containing 400 deformable cells. Cell pressure and global volume fraction of the cell volume is indicated. The capsule radius shrinks gradually so that equilibrium pressures are measured. The cell pressure may be slightly higher at the spheroid border due to arching effects of the outer cells.

**S4 Experimental Data. All_Experimental_data.xlsx** (sheet 1) provides the capsule data from [21] plus new data. Sheet 2 provides the dextran data that was extracted from [12].

## Footnotes

1 We assume *V/V_ref_* ≤ 1 in the experiment meaning the cells are always in a compressive state

2 The cell index has been dropped here for clarity.

